# ErbB dysregulation impairs cognition via myelination-dependent and - independent oligodendropathy

**DOI:** 10.1101/2020.09.09.289223

**Authors:** Xu Hu, Guanxiu Xiao, Li He, Qingyu Zhu, Xiaojie Niu, Huashun Li, Qi Xu, Zhengdong Wei, Hao Huang, Yifei Luan, Mengsheng Qiu, Kenji F. Tanaka, Ying Shen, Yanmei Tao

## Abstract

White matter abnormalities are an emerging pathological feature of schizophrenia. However, their attributions to the disease remain largely elusive. ErbB receptors and their ligands, some of which are essential for peripheral myelination, confer susceptibility to schizophrenia. By synergistically manipulating ErbB receptor activities in a oligodendrocyte-stage-specific manner in mice after early development, we demonstrate the distinct effects of ErbB signaling on oligodendrocytes at various differentiation states. ErbB overactivation, in mature oligodendrocytes, induces necroptosis causing demyelination, whereas in oligodendrocyte precursor cells, induces apoptosis causing hypomyelination. In contrast, ErbB inhibition increases oligodendrocyte precursor cell proliferation but induces hypomyelination by suppressing the myelinating capabilities of newly-formed oligodendrocytes. Remarkably, ErbB inhibition in mature oligodendrocytes diminishes axonal conduction under energy stress and impairs working memory capacity independently of myelin pathology. This study reveals the etiological implications of oligodendrocyte vulnerability induced by ErbB dysregulation, and elucidates the pathogenetic mechanisms for variable structural and functional white matter abnormalities.

## Introduction

Adolescence is the critical period for the central nervous system (CNS) to completely develop and mature. In particular, CNS myelin generated by oligodendrocytes (OLs) is one of the most developmentally active component in the adolescent brain. This may lead to CNS myelin being a highly susceptible target in psychiatric disorders such as schizophrenia which typically develops during adolescence (*Fields, 2008; Hoistad et al., 2009; Kessler et al., 2007; Peters and Karlsgodt, 2015*). A growing body of literature points to abnormalities in the structure, component proteins, or regulating molecules of CNS myelin in schizophrenic patients (*Douaud et al., 2007; Fields, 2008; Hof and Schmitz, 2009; Hoistad et al., 2009; Kelly et al., 2018; Uranova et al., 2011*). Schizophrenia is increasingly viewed as a spectrum disorder based on varied symptom severity and genetic risk. Especially, white-matter microstructural changes as examined by structural brain imaging techniques are sensitive to the symptom severity or genetic loading of schizophrenic patients (*Karlsgodt, 2020*). Therefore, understanding schizophrenia related myelin pathogenesis is crucial for the development of diagnostic standards or therapeutic targets given that it is one of the most promising features whose progression can be examined periodically in patients.

Tyrosine kinase receptors ErbB(1-4) mediate the signaling of numerous growth factors which are categorized into the neuregulin (NRG) family and the epidermal growth factor (EGF) family (*Iwakura and Nawa, 2013; Mei and Nave, 2014*). The NRG and EGF family members bind differentially to the four ErbB receptors. Due to the indispensable function of NRG1-ErbB signaling in peripheral myelination (*Nave and Salzer, 2006*), it was expected that NRG-ErbB signaling played a role in CNS myelin development. However, the contradictory results from different research groups have silenced any significance of this previous postulate (*Brinkmann et al., 2008; Makinodan et al., 2012; Schmucker et al., 2003; Taveggia et al., 2008*). Genetic ablation of *NRG1* or *ErbB4*, the ligand and receptor that have received extensive attention from researchers in schizophrenia field (*Harrison and Law, 2006; Mei and Nave, 2014*), induces neither developmental alteration nor pathogenesis in white matter of mutant mice (*Brinkmann et al., 2008*). However, studies combining genetic linkage analysis and brain imaging techniques have associated *NRG1* or *ERBB4* variability with reduced white matter density or integrity in human subjects (*McIntosh et al., 2008; Winterer et al., 2008; Zuliani et al., 2011*).

Notably, in addition to NRG1 and ErbB4, many molecules in the ErbB signaling pathways exhibit single nucleotide polymorphisms (SNPs) or aberrant expression that are implicated in schizophrenia or other psychiatric disorders. Both gain and loss of ErbB signaling have been indicated by genetic and biochemical studies (*Harrison and Law, 2006; Mei and Nave, 2014*). Particularly, NRG1 and ErbB4 have been revealed to increase the mRNA levels, protein levels, or receptor activity in the schizophrenic brain (*Chong et al., 2008; Hahn et al., 2006; Joshi et al., 2014; Law et al., 2006; Law et al., 2012*). It is noteworthy that EGFR (ErbB1), which only binds the EGF family ligands, is elevated in the brain of schizophrenic patients (*Futamura et al., 2002*) and shows potential in regulating oligodendrogenesis in developmental and pathological conditions (*Aguirre et al., 2007*). Thus, NRG-ErbB and EGF-ErbB signaling may be synergistic in the regulation of CNS myelin integrity.

In the CNS, OL precursor cells (OPCs) after terminal mitosis differentiate into newly-formed OLs (NFOs). NFOs then span differentiation states from pre-myelinating OLs to newly myelinating OLs. Myelinating OLs effectively generate myelin sheaths in a short time window before further differentiating into mature OLs (MOs) that maintain the myelin sheath (*Bergles and Richardson, 2015; Czopka et al., 2013; Hughes et al., 2018; Tripathi et al., 2017; Watkins et al., 2008; Xiao et al., 2016*). To study whether ErbB receptors, through mediating NRG-ErbB and EGF-ErbB signaling, cooperate to regulate OLs and CNS myelin, we adopted an inducible pan-ErbB strategy and manipulated ErbB receptor activities specifically in OL lineage cells *in vivo*. This strategy allowed us to avoid characterizing the complex composition of ErbB ligands or receptors in OL lineage cells and helped us focus on their cellular function. The results reveal that ErbB dysregulation differentially affects OPCs, NFOs, and MOs, leading to CNS demyelination, hypomyelination, and even OL dysfunction that causes cognitive deficits independently of myelin pathology.

## Results

### ErbB overactivation in OLs induces demyelination and hypomyelination

Studies on OL-specific knock-out mice have validated the expression of ErbB3 and ErbB4 in OLs (*Brinkmann et al., 2008*), while phosphorylated EGFR is detected in OLs by immunostaining (*Palazuelos et al., 2014*). We characterized the expression of ErbB receptor members in subcortical white matter regions at different postnatal days. Note that the subcortical white matter regions isolated from mice before P5 had few myelin components. Our results indicate that ErbB2 is barely expressed in mouse CNS myelin, whereas EGFR, ErbB3 and ErbB4 are expressed with relatively stable levels during juvenile to adolescent development (Figure 1-figure supplement 1A,B).

**Figure 1.**
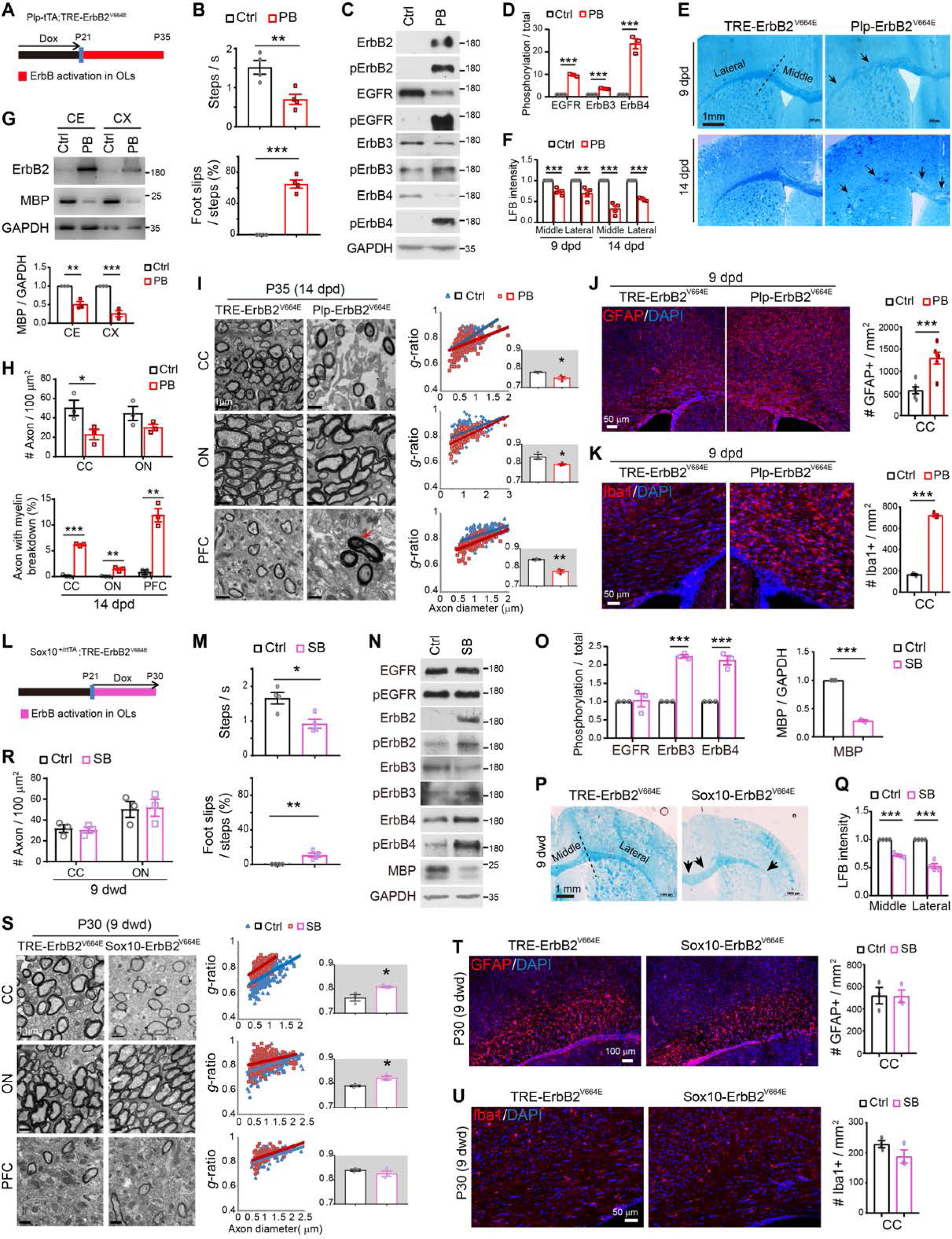
ErbB overactivation in OLs induces demyelination in *Plp*-ErbB2^V664E^ (PB) mice but hypomyelination in *Sox10*-ErbB2^V664E^ (SB) mice. (**A** and **L**) Dox treatment setting for indicated mice and littermate controls. (**B** and **M**) Walking speed and percentage of foot slips of *Plp*-ErbB2^V664E^ mice and littermate controls (Ctrl) at P35 with 14 dpd, or *Sox10*-ErbB2^V664E^ mice and littermate controls at 9 dwd, in the grid walking test. Data were presented as mean ± s.e.m., and analyzed by two-tailed unpaired *t* test. n = 4 mice for each group. In B, for velocity, *t*_(6)_ = 3.773, *P* = 0.0093; For foot slips, *t*_(6)_ = 12.31, *P* < 0.0001. In M, for velocity, *t*_(6)_ = 3.504, *P* = 0.0128; For foot slips, *t*_(6)_ = 4.429, *P* = 0.0044. (**C** and **N**) Western blotting of indicated proteins in white matter tissues isolated from *Plp*-ErbB2^V664E^ mice, or *Sox10*-ErbB2^V664E^ mice, in comparison with that from littermate control mice. Activation status of each ErbB receptor was examined by western blotting with the specific antibody against its phosphorylated form. (**D** and **O**) Quantitative data of western blotting results were presented as mean ± s.e.m., and analyzed by two-tailed unpaired *t* test. In D, for EGFR, *t*_(4)_ = 27.64, *P* < 0.0001; for ErbB3, *t*_(4)_ = 19.98, *P* < 0.0001; for ErbB4, *t*_(4)_ = 10.06, *P* = 0.0005. In O, for EGFR, *t*_(4)_ = 0.1983, *P* = 0.852; for ErbB3, *t*_(4)_ = 28.34, *P* < 0.0001; for ErbB4, *t*_(4)_ = 9.181, *P* = 0.00078; for MBP, *t*_(4)_ = 48.82, *P* < 0.0001. (**E** and **P**) LFB staining results of coronal sections through the genu of the corpus callosum in *Plp*-ErbB2^V664E^ and littermate control mice, or in *Sox10*-ErbB2^V664E^ and littermate control mice. Black arrows indicate the lower staining intensity of myelin stained in the corpus callosum. (**F** and **Q**) Quantitative data for LFB staining intensity in the corpus callosum of indicated mice. Data were presented as mean ± s.e.m. and analyzed by two-tailed unpaired *t* test. In F, for the middle part at 9 dpd, *t*_(6)_ = 6.345, *P* = 0.00072; for the lateral part at 9 dpd, *t*_(6)_ = 3.914, *P* = 0.0079; for the middle part at 14 dpd, *t*_(6)_ = 9.89, *P* < 0.0001; for the lateral part at 14 dpd, *t*_(6)_ = 23.07, *P* < 0.0001. In Q, for the middle part, *t*_(6)_ = 15.17, *P* < 0.0001; for the lateral part, *t*_(6)_ = 10.23, *P* < 0.0001. (**G**) Western blotting results of MBP and ErbB2 in the cortex (CX) and the cerebellum (CE) isolated from *Plp*-ErbB2^V664E^ and littermate control mice at 14 dpd. GAPDH served as loading control. Quantitative data were presented as mean ± s.e.m., and analyzed by two-tailed unpaired *t* test. For CE, *t*_(4)_ = 6.35, *P* = 0.0032; for CX, *t*_(4)_ = 9.243, *P* = 0.00076. (**H** and **R**) The densities of myelinated axons examined by EM in different brain regions of *Plp*-ErbB2^V664E^ and littermate control mice at 14 dpd, or *Sox10*-ErbB2^V664E^ and littermate control mice at 9 dwd, were quantified. The percentages of axons with myelin breakdown were also quantified for *Plp*-ErbB2^V664E^ mice (H). CC, the corpus callosum; ON, the optic nerve; PFC, the prefrontal cortex. Data were presented as mean ± s.e.m., and analyzed by two-tailed unpaired *t* test. In H, for myelinated-axon density, in CC, *t*_(4)_ = 2.863, *P* = 0.046; in ON, *t*_(4)_ = 1.818, *P* = 0.143. For the percentage of axons with myelin breakdown, in CC, *t*_(4)_ = 29.32, *P* < 0.0001; in ON, *t*_(4)_ = 6.108, *P* = 0.0036; in PFC, *t*_(4)_ = 8.125, *P* = 0.0012. In R, for myelinated-axon density, in CC, t_(4)_ = 0.2773, *P* = 0.795; in ON, t_(4)_ = 0.1455, *P* = 0.891. (**I** and **S**) EM images of the corpus callosum (CC), optic nerve (ON), and prefrontal cortex (PFC) from *Plp*-ErbB2^V664E^ and littermate controls at 14 dpd, or from *Sox10*-ErbB2^V664E^ and littermate control mice at 9 dwd. Red arrow indicates the axon with myelin breakdown. Quantitative data were shown for *g*-ratio analysis of myelinated axons detected by EM. Averaged g-ratio for each mouse were plotted as inset, presented as mean ± s.e.m., and analyzed by two-tailed unpaired *t* test. In I, for CC, *t*_(4)_ = 3.412, *P* = 0.027; for ON, *t*_(4)_ = 3.083, *P* = 0.037; for PFC, *t*_(4)_ = 7.11, *P* = 0.0021. In S, for CC, *t*_(4)_ = 3.295, *P* = 0.03; for ON, *t*_(4)_ = 3.775, *P* = 0.0195; for PFC, *t*_(4)_ = 1.196, *P* = 0.298. (**J**, **K**, **T**, **U**) Astrocytes (GFAP^+^) and microglia (Iba1^+^) examined in the subcortical white matter of indicated mice by immunostaining. Cell densities in the corpus callosum were quantified, and data were presented as mean ± s.e.m., and analyzed by two-tailed unpaired *t* test. In J, *t*_(10)_ = 4.753, *P* = 0.0008. In K, *t*_(4)_ = 36.4, *P* < 0.0001. In T, *t*_(4)_ = 0.0501, *P* = 0.962. In U, *t*_(4)_ = 1.637, *P* = 0.177.

To manipulate ErbB receptor activities in the OL lineage of mice, we employed tetracycline-controlled systems whose induction or blockade depends on the presence of doxycycline (Dox). We first characterized the impact of elevated ErbB receptor activities on CNS myelin by generation of *Plp*-tTA;*TRE*-ErbB2^V664E^ (*Plp*-ErbB2^V664E^) bi-transgenic ‘Tet-Off’ mice (Figure 1A). Notably, *Plp*-ErbB2^V664E^ mice around P35 exhibited severe ataxia while walking on a grid panel (Figure 1B). Moreover, *Plp*-ErbB2^V664E^ mice showed difficulty in rolling over, indicating severely impaired motor coordination.

With the expression and phosphorylation of ectopic ErbB2^V664E^, endogenous ErbB receptors (EGFR, ErbB3 and ErbB4) were strikingly phosphorylated and activated in the white matter of *Plp*-ErbB2^V664E^ mice (Figure 1C,D). Overactivation of ErbB receptors caused lower myelin staining intensity as exhibited in the corpus callosum of *Plp*-ErbB2^V664E^ mice at 9 days post Dox-withdrawal (dpd) after LFB staining. Notably, at 14 dpd, myelin loss became more evident throughout the corpus callosum, suggesting that *Plp*-ErbB2^V664E^ mice were undergoing CNS demyelination after Dox withdrawal (Figure 1E,F). Consistently, western blotting revealed loss of myelin basic protein (MBP), an indicator for MOs and myelin, in the brain of *Plp*-ErbB2^V664E^ mice (Figure 1G). Moreover, the electron microscopic examination (EM) of the myelin ultrastructure revealed that myelin sheaths of some axons in *Plp*-ErbB2^V664E^ mice ruptured or underwent breakdown (Figure 1H,I and Figure 1-figure supplement 2A), consistent with the idea of demyelination. Due to demyelination, only a few intact axons were detected in the midline of the corpus callosum of *Plp*-ErbB2^V664E^ mice at 14 dpd (Figure 1H,I). When axonal tracts in the corpus callosum were immunostained by TuJ1, the antibody recognizing neuronal specific β-tubulin III, the immunoreactivity dramatically reduced in *Plp*-ErbB2^V664E^ mice at 14 dpd (Figure 1-figure supplement 2B). In addition, as a pathological condition, demyelination is usually complicated and aggravated by the pathological responses from nearby astrocytes and microglia. Indeed, in the white matter of *Plp*-ErbB2^V664E^ mice, astrogliosis and microgliosis were revealed (Figure 1J,K).

Interestingly, despite the demyelination, the detectable axons, throughout the corpus callosum, optic nerve, and prefrontal cortex in *Plp*-ErbB2^V664E^ mice, were hypermyelinated. Myelinated axons detected in the brain of *Plp*-ErbB2^V664E^ mice had significantly smaller *g*-ratio (axon diameter/fiber diameter), a quantitative indication of myelin thickness for individual axons with different diameters (Figure 1I). Myelin thickness showed no difference between *TRE*-ErbB2^V664E^ and littermate *Plp*-tTA mice after Dox withdrawal (Figure 1-figure supplement 2C). Therefore, hypermyelination of detectable axons in *Plp*-ErbB2^V664E^ mice was a result of the overexpression of ErbB2^V664E^, which was detected by an antibody against ErbB2 (Figure 1C,G).

Hypermyelination of individual axons in *Plp*-ErbB2^V664E^ mice phenocopied that observed in NRG1-overexpressing mice (*Brinkmann et al., 2008*). Notably, EM examination of the ultrastructure in *Plp*-ErbB2^V664E^ mice at 9 dpd revealed that most axons were intact in the midline of the corpus callosum, although they have been significantly hypermyelinated (Figure 1-figure supplement 2D,E). These results confirmed that hypermyelination occurs early in *Plp*-ErbB2^V664E^ mice, and demyelination and axonal degeneration in *Plp*-ErbB2^V664E^ mice are pathological events induced secondarily by continuous ErbB activation.

Next, we examined the effects of ErbB activation on CNS myelin by a ‘Tet-on’ system generated in *Sox10*^+/rtTA^;*TRE*-ErbB2^V664E^ (*Sox10*-ErbB2^V664E^) mice (Figure 1L). *Sox10*-ErbB2^V664E^ mice with Dox feeding from P21 developed severe motor dysfunction, including ataxia and tremors, and died around P35. As a result, *Sox10*-ErbB2^V664E^ and littermate control mice were investigated at P30 after 9 days with Dox-feeding (dwd). These mice had smaller body sizes at P30 and walked with difficulty on a grid panel (Figure 1M). Western blotting revealed the expression and phosphorylation of ectopic ErbB2^V664E^ accompanied with increases in phosphorylation of ErbB3 and ErbB4, but not that of EGFR, in the white matter of *Sox10*-ErbB2^V664E^ mice (Figure 1N,O). Brain slices stained by LFB exhibited lower staining intensity in the white matter of *Sox10*-ErbB2^V664E^ mice (Figure 1P,Q), consistent with the lower MBP levels detected by western blotting (Figure 1N,O). The examination of the ultrastructure by EM revealed that the axons in the corpus callosum and optic nerve of *Sox10*-ErbB2^V664E^ mice exhibited thinner myelin with significantly increased *g*-ratio (Figure 1S). *Sox10*^+/rtTA^ is a knock-in mouse line, so that the allele with *Sox10*-rtTA would not transcribe *Sox10* mRNA (*Ludwig et al., 2004*). We analyzed the ultrastructure of myelinated axons in *Sox10*^+/rtTA^ and littermate *TRE*-ErbB2^V664E^ mice at P30 and did not observe any differences (Figure 1-figure supplement 2F), therefore we can exclude the possible effect of haploinsufficiency of Sox10 on late postnatal myelin development.

It is notable that in *Sox10*-ErbB2^V664E^ mice, the numbers of myelinated axons were not altered (Figure 1R), and myelin sheaths exhibited normal morphology (Figure 1S). Moreover, neither microgliosis nor astrogliosis was detected in the white matter of *Sox10*-ErbB2^V664E^ mice (Figure 1T,U). Because there was no indication of inflammatory pathogenesis, we can conclude that thinner myelin in *Sox10*-ErbB2^V664E^ white matter are caused by developmental deficits not pathological conditions. Therefore, ErbB activation in *Sox10*-ErbB2^V664E^ mice induces CNS hypomyelination rather than demyelination.

### *Plp*-tTA targets mainly MOs whereas *Sox10*^+/rtTA^ targets OPC-NFOs

The finding that *Plp*-ErbB2^V664E^ and *Sox10*-ErbB2^V664E^ mice had no overlaps in histological and biochemical phenotypes was unexpected considering that Sox10 is reported to express throughout the OL lineage, and *Sox10*^+/rtTA^ knock-in mice have been used to investigate all OL lineage cells (*Wegener et al., 2015*). Because the induction of the ‘Tet-on’ or ‘Tet-off’ system by Dox feeding or Dox withdrawal has a delayed effect on gene expression, and reporter proteins could accumulate to label the consecutive cellular stages, the results obtained by using *TRE*-controlled reporter mice fail to accurately reveal the original cells targeted by tTA or rtTA. To circumvent this problem, we delivered a *TRE*-controlled fluorescence reporter carried by an adeno-associated virus (AAV) into the mouse brains at P14 or P35, and examined tTA- or rtTA-targeted cells as well as their derivatives within 2 days.

*Plp*-tTA mice were raised with no Dox feeding, whereas *Sox10*^+/rtTA^ were fed with Dox for 3 days before the stereotactic injection (Figure 2A). One or 2 days after virus injection, the reporter-expressing (YFP^+^) cells were all immunopositive for Olig2 in both mouse lines at either age (Figure 2-figure supplement 1A-D), confirming their OL lineage specificity. To analyze the differentiation stage specificity, we immunostained AAV-*TRE*-YFP infected brain slices with an antibody for NG2 or PDGFRα that labels OPCs, or the antibody CC1 that labels post-mitotic OLs. The results showed that very few (4-7%) YFP^+^ cells were OPCs, while 92-97% of them were post-mitotic OLs 1 or 2 days after virus injection in *Plp*-tTA mice at either age, as well as in *Sox10*^+/rtTA^ mice at P14 (Figure 2B,C and Figure 2-figure supplement 1A-D). However, for *Sox10*^+/rtTA^ mice at P35, approximately 26% of YFP^+^ cells were OPCs 1 day after virus injection, and it decreased to 8% after 1 more day (Figure 2C). It is known that OPCs can differentiate into NFOs as quick as 2.5 hours (*Xiao et al., 2016*). These results may suggest that most OPCs targeted by *Sox10*-rtTA at P35 are undergoing terminal differentiation (tOPCs), and *Sox10*-rtTA increasingly targets tOPCs from P14 to P35.

**Figure 2.**
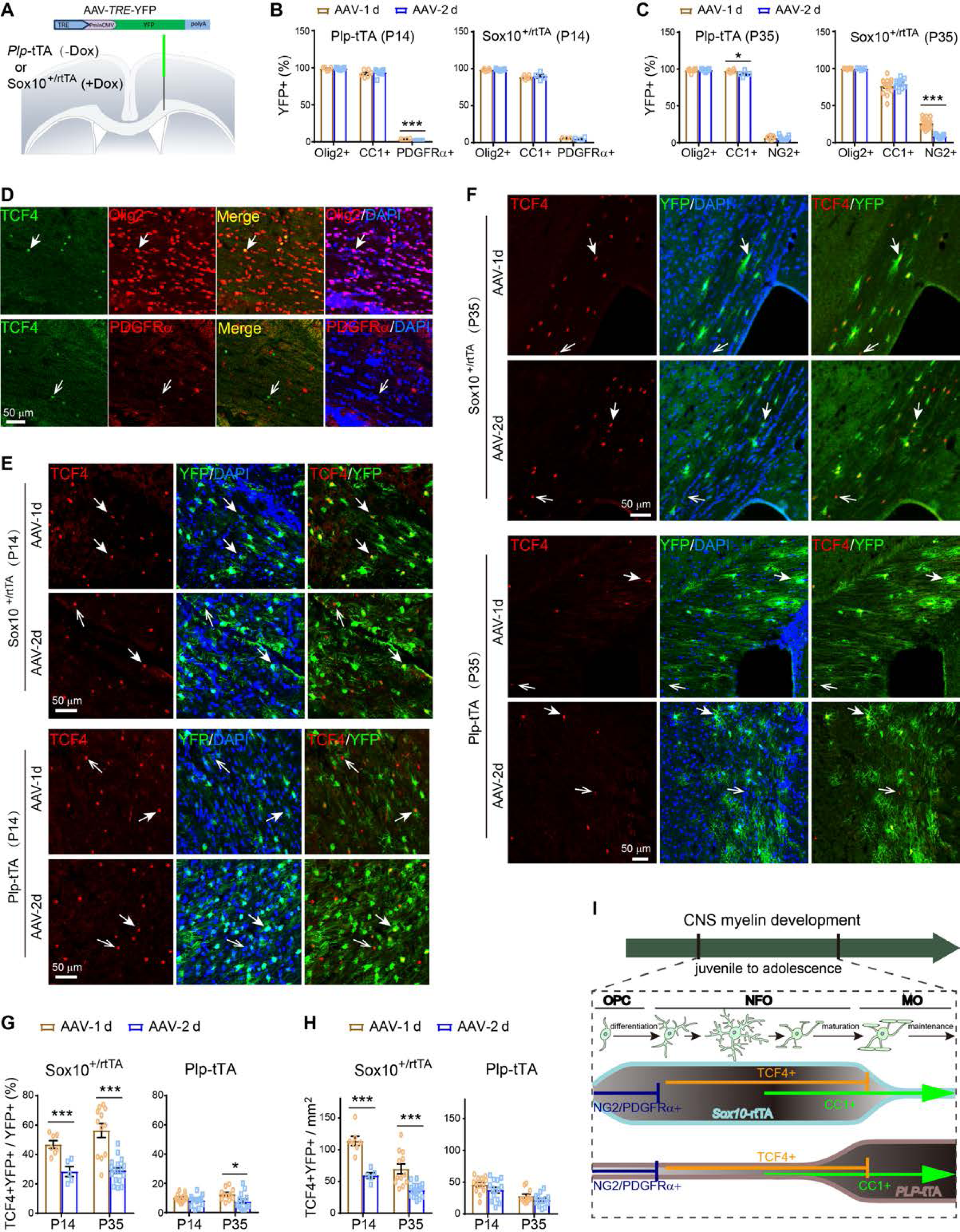
*Plp*-tTA targets MOs whereas *Sox10*^+/rtTA^ targets OPC-NFOs in mouse brains in late postnatal development. (**A**) Schematic illustration of stereotactic injection sites of AAV-*TRE*-YFP. (**B** and **C**) The percentage of Olig2^+^YFP^+^, CC1^+^YFP^+^, or PDGFRα^+^YFP^+^ (NG2^+^YFP^+^) cells in YFP^+^ cells for indicated mice 1 (AAV-1d) or 2 days (AAV-2d) after AAV-*TRE*-YFP injection at P14 (B) or P35 (C). Data were from repeated immunostaining results of 3-7 mice for each group, presented as mean ± s.e.m., and analyzed by two-tailed unpaired *t* test. For *Plp*-tTA at P14 from AAV-1d to AAV-2d: Olig2^+^YFP^+^ cells, *t*_(12)_ = 0.3698, *P* = 0.718; CC1^+^YFP^+^ cells, *t*_(13)_ = 0.5666, *P* = 0.581; PDGFRα^+^YFP^+^ cells, *t*_(10)_ = 7.532, *P* < 0.0001. For *Sox10*^+/rtTA^ at P14 from AAV-1d to AAV-2d: Olig2^+^YFP^+^ cells, *t*_(13)_ = 0.2055, *P* = 0.840; CC1^+^YFP^+^ cells, *t*_(8)_ = 0.6425, *P* = 0.539; PDGFRα^+^YFP^+^ cells, *t*_(5)_ = 1.021, *P* = 0.354. For *Plp*-tTA at P35 from AAV-1d to AAV-2d: Olig2^+^YFP^+^ cells, *t*_(17)_ = 0.4959, *P* = 0.626; CC1^+^YFP^+^ cells, *t*_(9)_ = 2.32, *P* = 0.046; NG2^+^YFP^+^ cells, *t*_(18)_ = 1.003, *P* = 0.329. For *Sox10*^+/rtTA^ at P35 from AAV-1d to AAV-2d: Olig2^+^YFP^+^ cells, *t*_(11)_ = 1.098, *P* = 0.296; CC1^+^YFP^+^ cells, *t*_(23)_ = 0.8614, *P* = 0.398; NG2^+^YFP^+^ cells, *t*_(26)_ = 7.869, *P* < 0.0001. (**D**) Immunostaining revealed that TCF4 was specifically expressed in a small fraction of Olig2^+^ cells, but not in PDGFRα^+^ cells, in the corpus callosum of mice at P30. Solid arrow, the representative double positive cell; Open arrow, the representative cell positive for TCF4 only. (**E** and **F**) Double immunostaining results of TCF4 and YFP for brain slices from indicated mice 1 (AAV-1d) or 2 days (AAV-2d) after virus injection at P14 (E) or P35 (F). Note that TCF4^+^ nuclei in AAV-infected area were almost all localized in YFP^+^ cells in *Sox10*^+/rtTA^ mice 1 day after virus injection (AAV-1d) at either P14 or P35, and the colocalization reduced after 1 more day (AAV-2d). Solid arrows, representative double positive cells; Open arrows, representative cells positive for TCF4 only. (**G** and **H**) The percentage of TCF4^+^YFP^+^ in YFP^+^ cells (G), and the density of TCF4^+^YFP^+^ cells (H), were analyzed. Data were from repeated immunostaining results of 3-7 mice for each group, presented as mean ± s.e.m., and analyzed by two- tailed unpaired *t* test. For the percentage in *Sox10*^+/rtTA^ mice from AAV-1d to AAV- 2d: at P14, *t*_(10)_ = 4.39, *P* = 0.0014; at P35, *t*_(28)_ = 6.041, *P* < 0.0001. For the percentage in *Plp*-tTA mice from AAV-1d to AAV-2d: at P14, *t*_(26)_ = 1.574, *P* = 0.128; at P35, *t*_(22)_ = 2.367, *P* = 0.027. For the density in *Sox10*^+/rtTA^ mice from AAV-1d to AAV-2d: at P14, *t*_(10)_ = 5.685, *P* = 0.0002; at P35, *t*_(28)_ = 4.813, *P* < 0.0001. For the density in *Plp*-tTA mice from AAV-1d to AAV-2d: at P14, *t*_(26)_ = 1.581, *P* = 0.126; at P35, *t*_(22)_ = 1.429, *P* = 0.167. (**I**) Schematic summary of OL stage-targeting preferences of *Plp*-tTA or *Sox10*^+/rtTA^ during juvenile to adolescent development.

There are also OL lineage cells belonging to the NFO stage that includes a transition from CC1^-^ to CC1^+^. β-catenin effector TCF4 is specifically expressed in the NFO stage (*Fancy et al., 2009; Fu et al., 2009; Ye et al., 2009*), which is present in a subset of Olig2^+^ cells but is absent in OPCs (PDGFRα^+^) in mice at P30 (Figure 2D). In *Sox10*^+/rtTA^ mice at P14, immunostaining revealed that approximately 49% of YFP^+^ cells were NFOs (TCF4^+^) 1 day after virus injection, but it reduced to 28% after 1 more day (Figure 2E,G). TCF4^+^ cells found in the corpus callosum at P35 were far fewer than P14, and these cells appeared as clusters (Figure 2-figure supplement 1E). Interestingly, in *Sox10*^+/rtTA^ mice at P35, YFP^+^ cells were mostly found in regions with TCF4^+^ cell clusters (Figure 2-figure supplement 1E), where approximately 56% of YFP^+^ cells were TCF4^+^ 1 day after virus injection and it reduced to 29% after 1 more day (Figure 2F,G). The half reduction of NFO percentage in YFP^+^ cells from day 1 to day 2 was consistent with the previous report that NFOs differentiate into MOs in 1 or 2 days (*Xiao et al., 2016*). There was another possibility that the transcriptional activity of *Sox10*-rtTA was low in MOs, and thus took more days to generate detectable YFP levels. We analyzed the densities of TCF4^+^YFP^+^ cells and found that they reduced to half from day 1 to day 2 after viral infection in *Sox10*^+/rtTA^ mice at either P14 or P35 (Figure 2H). The results excluded the possibility that the reduction of NFO ratio in YFP^+^ cells was due to the increase of targeted MOs, and confirmed the maturation of labeled NFOs from day 1 to day 2. A similar transition rate is applicable for targeted NFOs from day 0 to day 1. Therefore, these results corroborate that the majority of cells targeted by *Sox10*-rtTA at the time of AAV-*TRE*-YFP infection were tOPCs and NFOs (Figure 2I).

AAV-*TRE*-YFP in *Plp*-tTA mice also labeled some TCF4^+^ cells, which comprised only 7-12% of YFP^+^ cells 1 or 2 days after virus injection at either age (Figure 2E-H). It was noticeable that *Plp*-tTA did not specifically target TCF4^+^ cell clusters in the corpus callosum at P35 (Figure 2-figure supplement 1F). These results implied that *Plp*-tTA did not specifically target the tOPC or NFO stage but randomly expressed in the OPC-NFO period at a low ratio. Conversely, 90% of the YFP^+^ cells were TCF4^-^ and 92-97% were CC1^+^ in *Plp*-tTA mice at either P14 or P35, suggesting that *Plp*-tTA steadily targets MOs after early development (Figure 2I).

### ErbB overactivation causes MO necroptosis and OPC apoptosis

With an understanding of the differentiation stage-specific targeting preferences of *Sox10*^+/rtTA^ and *Plp*-tTA mice, we investigated the cellular mechanisms that determine the different myelin responses in *Plp*-ErbB2^V664E^ and *Sox10*-ErbB2^V664E^ mice. We found intact post-mitotic OLs (CC1^+^) decreased in the corpus callosum of *Plp*-ErbB2^V664E^ mice starting from 6 dpd (Figure 3A-C). The CC1^+^ cells with striking number reduction were MOs because the densities of NFOs (TCF4^+^) did not show significant change (Figure 3-figure supplement 1B,C). Meanwhile, OPCs (NG2^+^Olig2^+^) pathologically regenerated (Ki67^+^Olig2^+^) in the trunk of corpus callosum, indicating that the pathogenetic factor was released in the myelin-enriched region (Figure 3-figure supplement 1A-E).

**Figure 3.**
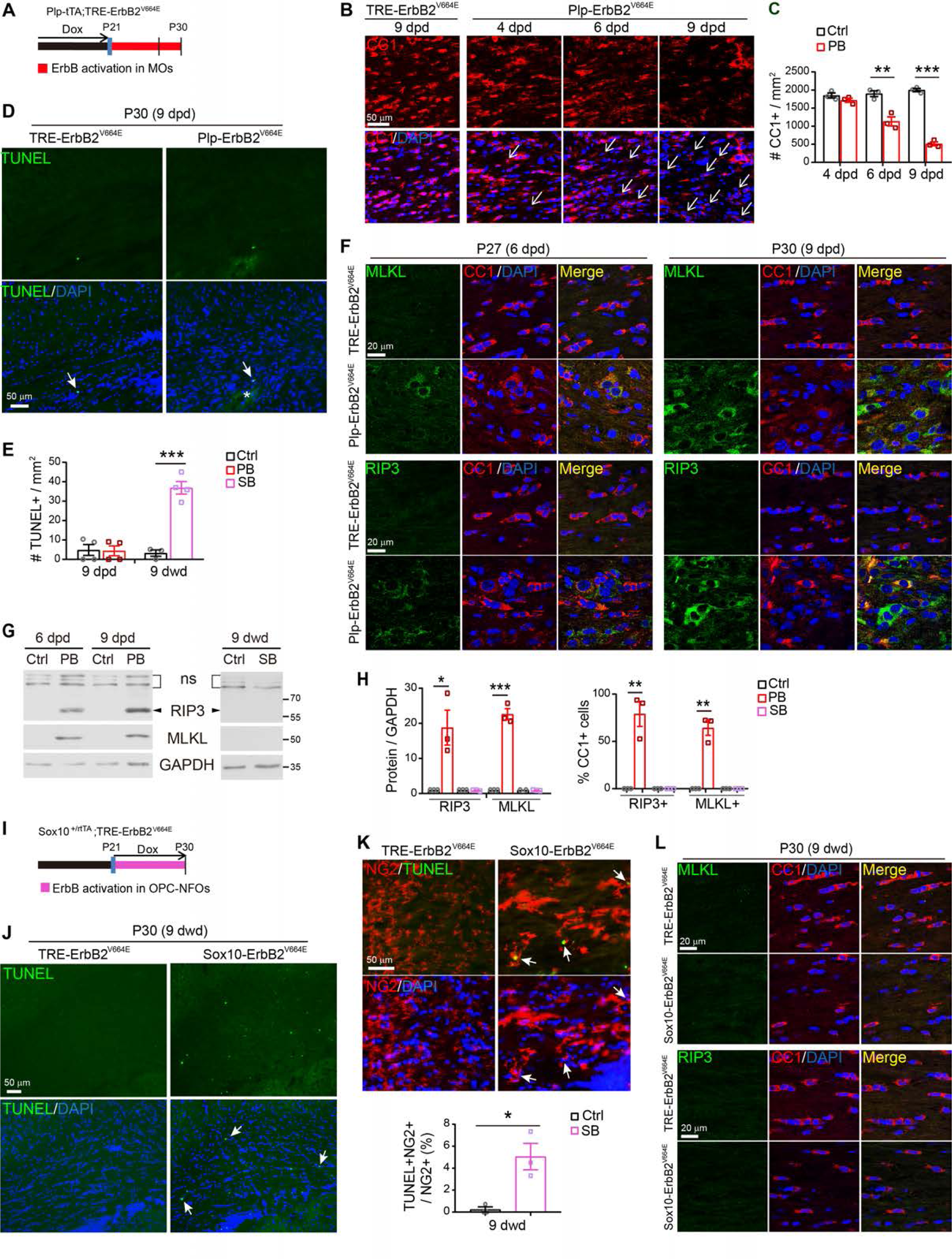
ErbB overactivation induces MO necroptosis in *Plp*-ErbB2^V664E^ (PB) mice but OPC apoptosis in *Sox10*-ErbB2^V664E^ (SB) mice. (**A** and **I**) Dox treatment setting for indicated mice and littermate controls. (**B**) The numbers of degenerating OLs (represented by nuclei associated with CC1^+^ cell debris, white arrows) increased in the corpus callosum of *Plp*-ErbB2^V664E^ mice starting from 6 dpd as revealed by CC1 immunostaining. (**C**) Quantitative data of intact CC1^+^ density in *Plp*-ErbB2^V664E^ and control mice (Ctrl) with indicated Dox treatment were present as mean ± s.e.m., and analyzed by two-tailed unpaired *t* test. For 4 dpd, *t*_(4)_ = 1.485, *P* = 0.212; for 6 dpd, *t*_(4)_ = 5.203, *P* = 0.0065; for 9 dpd, *t*_(4)_ = 20.95, *P* < 0.0001. (**D** and **J**) Apoptotic cells (arrows) in the corpus callosum of *Plp*-ErbB2^V664E^ and control mice at 9 dpd (D), or *Sox10*-ErbB2^V664E^ and control mice at 9 dwd (J), were examined by TUNEL assays. Note the increased numbers of nuclei (DAPI^+^) and non-specifically stained hemorrhagic spot (asterisk), which is the consequence of inflammation, in the brain slice of *Plp*-ErbB2^V664E^ mice (D). (**E**) Quantitative data of apoptotic cell densities in indicated mice. Data were presented as mean ± s.e.m., and analyzed by two-tailed unpaired *t* test. For PB, *t*_(6)_ = 0.1128, *P* = 0.914; for SB, *t*_(5)_ = 8.344, *P* = 0.0004. (**F** and **L**) Co-immunostaining results of MLKL or RIP3 with CC1 in the corpus callosum of indicated mice with indicated Dox treatments. (**G**) Western blotting results of MLKL and RIP3 in the white matter of *Plp*-ErbB2^V664E^ mice, or in that of *Sox10*-ErbB2^V664E^ mice, and in those of littermate control mice. ns, non-specific bands. (**H**) Quantitative data of immunostaining and western blotting results of MLKL or RIP3 in indicated mice at 9 days with Dox treatment. Data were presented as mean ± s.e.m., and analyzed by two-tailed unpaired *t* test. In western blotting results, for RIP3 protein in PB, *t*_(4)_ = 3.579, *P* = 0.023; for MLKL protein in PB, *t*_(4)_ = 13.69, *P*= 0.00017. In immunostaining results, for percentage of RIP3^+^ in CC1^+^ cells in PB, *t*_(4)_ = 6.002, *P* = 0.0039; for percentage of MLKL^+^ in CC1^+^ cells in PB, *t*_(4)_ = 8.202, *P* = 0.0012. (**K**) Apoptotic cells (TUNEL^+^) were OPCs (NG2^+^) in *Sox10*-ErbB2^V664E^ mice at P30 with 9 dwd. Arrows, representative double positive cells. Note OPCs in *Sox10*-ErbB2^V664E^ mice were hypertrophic. The percentage of TUNEL^+^NG2^+^ cells in NG2^+^ cells were quantified and data were presented as mean ± s.e.m. and analyzed by two-tailed unpaired *t* test. *t*_(4)_ = 3.95, *P* = 0.0168.

MO number reduction occurred earlier than the time when demyelination was obviously observed in the corpus callosum of *Plp*-ErbB2^V664E^ mice, suggesting that oligodendropathy may be the cause of demyelination. We examined the corpus callosum of *Plp*-ErbB2^V664E^ mice by TdT-mediated dUTP nick end labeling (TUNEL) assay, and observed as little apoptotic signaling as that in controls (Figure 3D,E). This result reveals that the degenerating CC1^+^ cells were necrotic rather than apoptotic. Consistently, the OL nuclei associated with the destroyed myelin sheaths in *Plp*-ErbB2^V664E^ mice were regular nuclei without apoptotic chromatin condensation (Figure 1-figure supplement 2A). In support of this theory, MLKL, the protein mediating cell necroptosis (*Cai et al., 2014*) as well as the peripheral myelin breakdown after nerve injury (*Ying et al., 2018*), demonstrated an increased expression in the callosal CC1^+^ cells in *Plp*-ErbB2^V664E^ mice from 6 dpd (Figure 3F-H). Necroptosis is a programmed form of necrosis. RIP3 is the kinase at the upstream of MLKL in this programmed death signaling pathway (*Ofengeim et al., 2015; Sun et al., 2012*). Notably, the expression of RIP3 was also elevated in the callosal CC1^+^ cells in *Plp*-ErbB2^V664E^ mice from 6 dpd as revealed by both immunostaining and western blotting (Figure 3F-H). Based on the timeline, MO necroptosis was the primary defect induced in *Plp*-ErbB2^V664E^ mice, followed by myelin breakdown, OPC regeneration, axonal degeneration, and other pathological events as reported in multiple sclerosis (*Bradl and Lassmann, 2010; Ofengeim et al., 2015*).

In contrast, for *Sox10*-ErbB2^V664E^ mice, a dramatic increase in cell apoptosis (TUNEL^+^) in the corpus callosum was observed (Figure 3E,I,J). These apoptotic nuclei were localized in NG2^+^ cells (Figure 3K), indicating apoptosis of OPCs. On the other hand, no increase of MLKL or RIP3 was detected (Figure 3G,H,L), indicating there was no necroptosis. Notably, both the NG2^+^ cells with and without TUNEL^+^ nuclei were hypertrophic in *Sox10*-ErbB2^V664E^ mice (Figure 3K). This phenomenon was not revealed for NG2^+^ cells in *Plp*-ErbB2^V664E^ mice (Figure 3-figure supplement 1B).

### ErbB inhibition in OPC-NFOs, but not in MOs, induces hypomyelination

Next, to investigate whether ErbB receptors are functionally required for OLs, we examined the effects of inhibiting ErbB activities in different OL stages on CNS myelin. To this end, we first generated *Sox10*^+/rtTA^;*TRE*-dnEGFR (*Sox10*-dnEGFR) mice (Figure 4A and Figure 4-figure supplement 1A,B). In line with the inhibitory effect of dnEGFR on endogenous ErbB activities (*Chen et al., 2017*), phosphorylation of ErbB3 and ErbB4 was reduced in white matter from *Sox10*-dnEGFR mice at P35 (Figure 4B,C). The phosphorylation of EGFR was not altered in *Sox10*-dnEGFR mice at P35 (Figure 4B,C), consistent with the finding in *Sox10*-ErbB2^V664E^ mice (Figure 1N,O). Myelin thickness and ultrastructures in the white matter of *Sox10*-dnEGFR and littermate control mice at P35 with 14 dwd did not show significant differences (Figure 4-figure supplement 1C). Therefore, we raised these mice to adulthood with continuous Dox feeding. Phosphorylation of EGFR, instead of ErbB3 or ErbB4, was apparently reduced in the white matter of *Sox10*-dnEGFR mice at P65 (Figure 4B,C). This change of ErbB receptors targeted by dnEGFR in *Sox10*-dnEGFR mice implied a switch of functional NRG-ErbB signaling to EGF-ErbB signaling in *Sox10*-rtTA-targeted cells from adolescence to adulthood.

**Figure 4.**
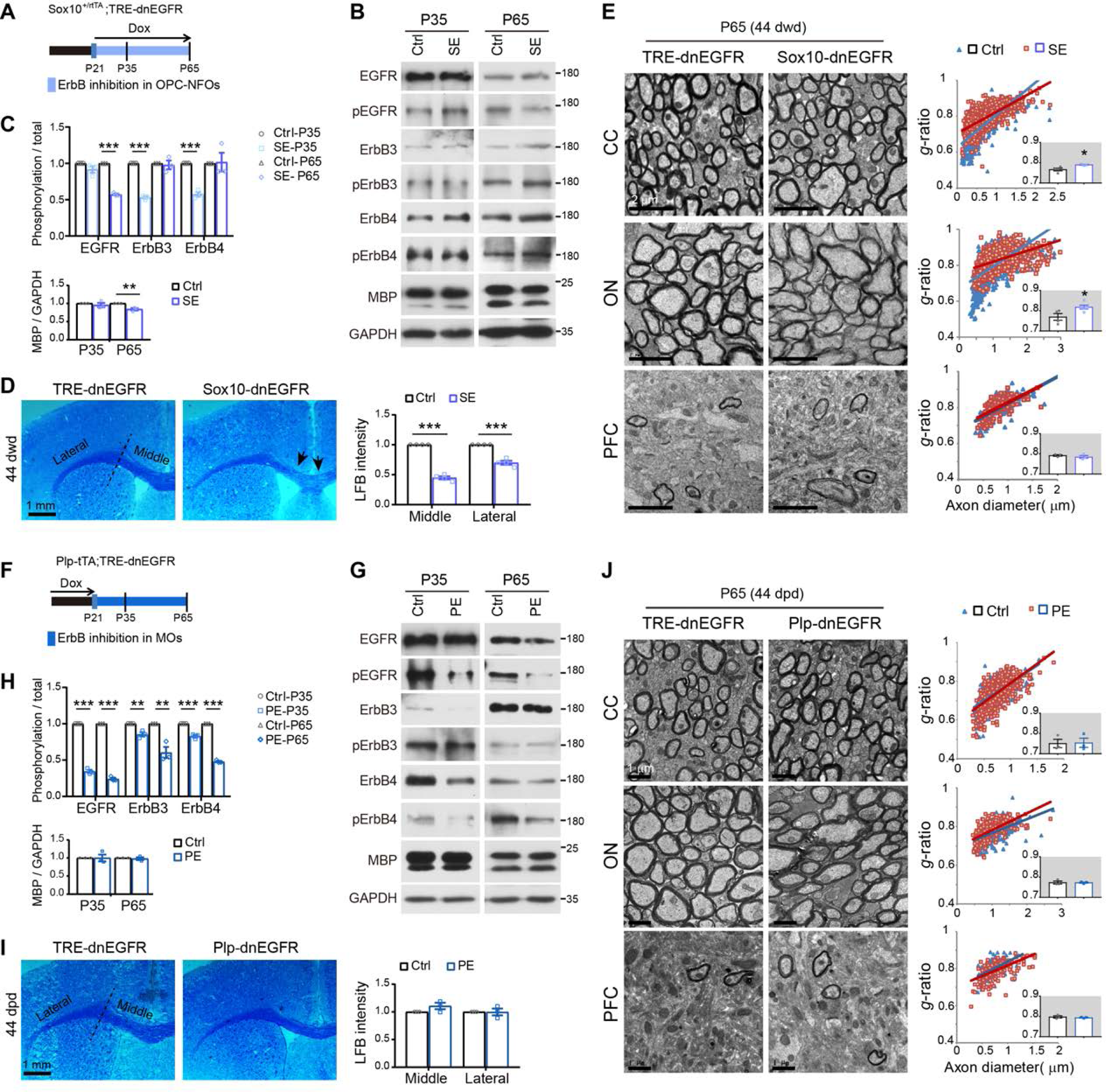
ErbB inhibition induces hypomyelination in *Sox10*-dnEGFR (SE) mice, but no myelin alteration in *Plp*-dnEGFR (PE) mice. (**A** and **F**) Dox treatment setting for indicated mice and littermate controls. (**B** and **G**) MBP levels and the inhibited phosphorylation in ErbB receptors were examined by western blotting in the white matter of *Sox10*-dnEGFR mice, or *Plp*-dnEGFR mice, in comparison with that of littermate controls (Ctrl). (**C** and **H**) Quantitative data of western blotting results. Data were presented as mean ± s.e.m., and analyzed by two-tailed unpaired *t* test. In C at P35, for EGFR, *t*_(4)_ = 1.813, *P* = 0.144; for ErbB3, *t*_(4)_ = 25.94, *P* < 0.0001; for ErbB4, *t*_(4)_ = 12.69, *P* = 0.00022; for MBP, *t*_(4)_ = 0.7711, *P* = 0.484. In C at P65, for EGFR, *t*_(4)_ = 35.09, *P* < 0.0001; for ErbB3, *t*_(4)_ = 0.3492, *P* = 0.745; for ErbB4, *t*_(4)_ = 0.138, *P* = 0.897; for MBP, *t*_(4)_ = 4.842, *P* = 0.0084. In H at P35, for EGFR, *t*_(4)_ = 28.36, *P* < 0.0001; for ErbB3, *t*_(4)_ = 4.925, *P* = 0.0079; for ErbB4, *t*_(4)_ = 8.838, *P* = 0.0009; for MBP, *t*_(4)_ = 0.00896, *P* = 0.993. In H at P65, for EGFR, *t*_(4)_ = 43.97, *P* < 0.0001; for ErbB3, *t*_(4)_ = 5.157, *P* = 0.0067; for ErbB4, *t*_(4)_ = 44.67, *P* < 0.0001; for MBP, *t*_(4)_ = 0.4686, *P* = 0.664. (**D** and **I**) LFB staining results of coronal sections through the genu of the corpus callosum in *Sox10*-dnEGFR and control mice at P65 with 44 dwd (D), or *Plp*-dnEGFR and control mice at P65 with 44 dpd (I). Black arrows indicate the middle part of the corpus callosum in *Sox10*-dnEGFR mice exhibiting obvious lower staining intensity. Quantitative data of LFB intensity were presented as mean ± s.e.m., and analyzed by two-tailed unpaired *t* test. In D, for the middle part, *t*_(6)_ = 21.18, *P* < 0.0001; for the lateral part, *t*_(6)_ = 9.121, *P* < 0.0001. In I, for the middle part, *t*_(4)_ = 1.814, *P* = 0.144; for the lateral part, *t*_(4)_ = 0.0287, *P* = 0.979. (**E** and **J**) EM images of the corpus callosum (CC), optic nerve (ON), and prefrontal cortex (PFC) from *Sox10*-dnEGFR and littermate controls at 44 dwd (E), or *Plp*-dnEGFR and littermate controls at 44 dpd (J). *g*-ratio was calculated for myelinated axons and averaged *g*-ratio were analyzed by two-tailed unpaired *t* test (inset). In E, for CC, *t*_(4)_ = 2.793, *P* = 0.0383; for ON, *t*_(7)_ = 2.629, *P* = 0.0339; for PFC, *t*_(4)_ = 0.8697, *P* = 0.434. In J, for CC, *t*_(4)_ = 0.1139, *P* = 0.915; for ON, *t*_(4)_ = 0.0754, *P* = 0.944; for PFC, *t*_(4)_ = 0.6334, *P* = 0.561.

For *Sox10*-dnEGFR mice at P65 with 44 dwd, myelin stained by LFB exhibited reduced intensity in the trunk of the corpus callosum (Figure 4D). Moreover, axons in the corpus callosum and optic nerve of *Sox10*-dnEGFR mice at P65 were hypomyelinated in comparison with that of littermate controls (Figure 4E). Consistently, MBP was reduced in the white matter of *Sox10*-dnEGFR mice at P65 (Figure 4B,C). Therefore, ErbB inhibition in OPC-NFOs starting from P21 results in hypomyelination in adulthood.

On the other hand, we crossed *Plp*-tTA and *TRE*-dnEGFR to generate *Plp*-tTA;*TRE*-dnEGFR (*Plp*-dnEGFR) mice (Figure 4F and Figure 4-figure supplement 1D,E). Western blotting revealed significant suppression on the phosphorylation of EGFR, as well as a mild suppression on that of ErbB3 and ErbB4 (Figure 4G,H), consistent with their overactivation in *Plp*-ErbB2^V664E^ mice (Figure 1C,D). No CNS myelin differences were observed in *Plp*-dnEGFR and littermate control mice at P35 with 14 dpd (Figure 4-figure supplement 1F).

We extended our investigation to P65 when dnEGFR still functionally suppressed ErbB receptor activities in the white matter of *Plp*-dnEGFR mice (Figure 4G,H). Even in the adult mice, when dnEGFR had been expressed in MOs for 44 days, the brains of *Plp*-dnEGFR and littermate mice exhibited no difference in LFB-stained myelin (Figure 4I), MBP protein levels (Figure 4G,H), or myelin ultrastructures (Figure 4J). These results suggest that the dual blockade of endogenous NRG-ErbB and EGF-ErbB signaling in MOs does not affect CNS myelin integrity, and that ErbB activities are not required for the maintenance of CNS myelin after maturation.

### ErbB activation blocks OPC proliferation and survival, whereas promotes NFO differentiation and myelination

It is intriguing to note that both ErbB inhibition and overactivation in OPC-NFOs result in hypomyelination in the CNS. This implies that ErbB signaling can regulate CNS myelin development by different mechanisms. To investigate the mechanisms, we first compared the states of OL lineage cells in *Sox10*-ErbB2^V664E^ and *Sox10*-dnEGFR mice. In line with the finding that OPCs underwent apoptosis, the numbers of Olig2^+^, NG2^+^, and CC1^+^ cells significantly decreased in the corpus callosum of *Sox10*-ErbB2^V664E^ mice (Figure 5A and Figure 5-figure supplement 1A), while there was no OPC or OL differences between *Sox10*^+/rtTA^ and *TRE*-ErbB2^V664E^ littermates at P30 with 9 dwd (Figure 5-figure supplement 1A). Further, we found proliferating OPCs (Ki67^+^Olig2^+^) significantly decreased in the corpus callosum of *Sox10*-ErbB2^V664E^ mice (Figure 5D,G). In contrast, the densities of proliferating OPCs (Ki67^+^Olig2^+^) and Olig2^+^ cells increased in *Sox10*-dnEGFR mice at P65 (Figure 5B,E,G and Figure 5-figure supplement 1B), despite the fact that these increases were not observed in *Sox10*-dnEGFR mice at P35 (Figure 5-figure supplement 2A-C). No pathological signs were observed and the increased Olig2^+^ cells comprised of both NG2^+^ and CC1^+^ cells (Figure 5B and Figure 5-figure supplement 1B). It could not be determined whether apoptosis decreased in *Sox10*-dnEGFR white matter, as the apoptotic cells (TUNEL^+^) were minimal in white matter of both *Sox10*-dnEGFR and control mice (Figure 5H). The consistent results from gain-of-function (*Sox10*-ErbB2^V664E^) and loss-of-function (*Sox10*-dnEGFR) studies support a negative regulation of OPC proliferation and survival by ErbB signaling.

**Figure 5.**
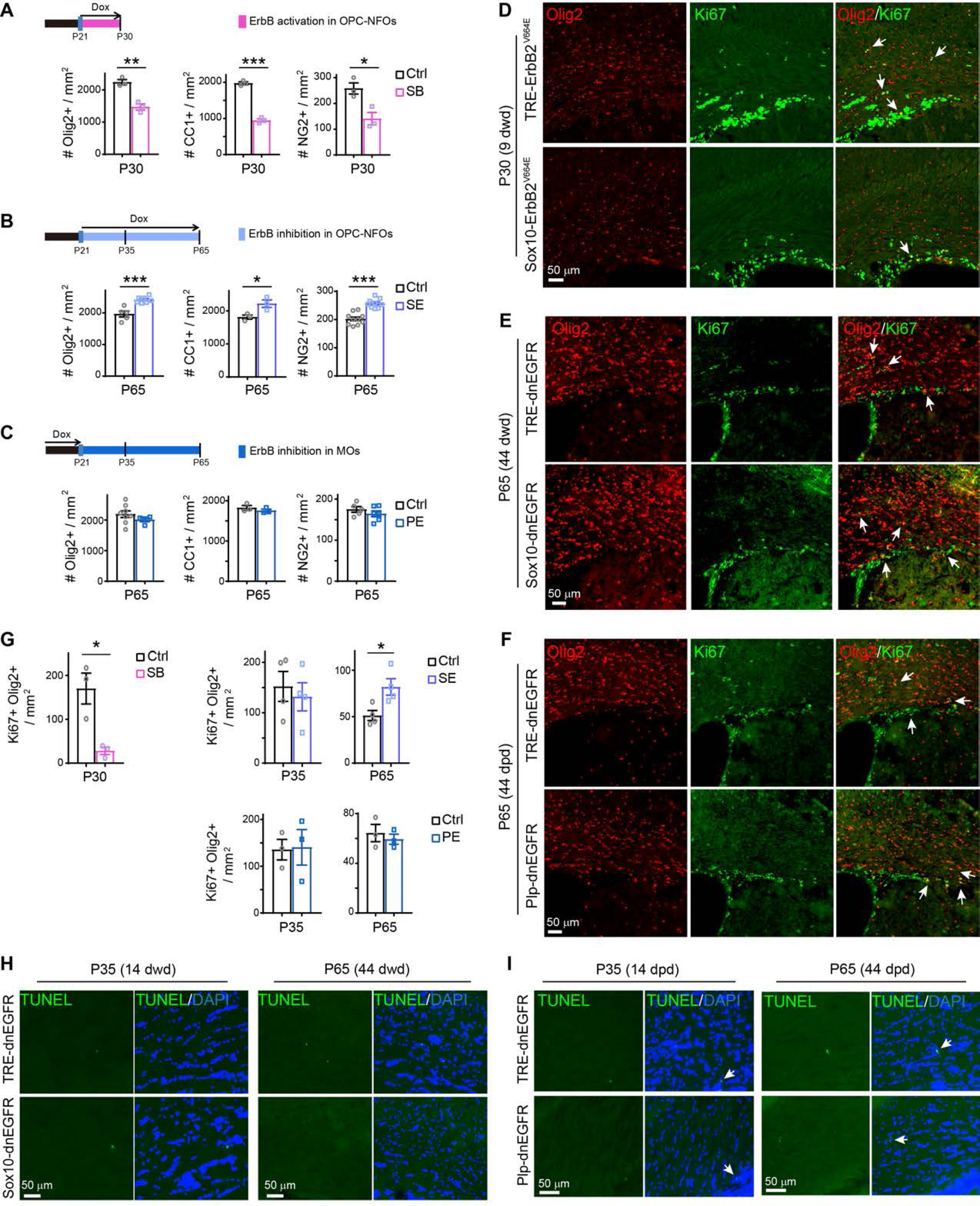
ErbB activation negatively regulates OPC proliferation. (**A**-**C**) Statistic results of Olig2^+^, CC1^+^, and NG2^+^ cell densities in the corpus callosum of *Sox10*-ErbB2^V664E^ mice (SB) and littermate controls (Ctrl) at P30 with 9 dwd, *Sox10*-dnEGFR mice (SE) and littermate controls at P65 with 44 dwd, or *Plp*-dnEGFR mice (PE) and littermate controls at P65 with 44 dpd. Data were from repeated immunostaining of 3 mice for each group, presented as mean ± s.e.m., and analyzed by two-tailed unpaired *t* test. In A, for Olig2^+^, *t*_(4)_ = 6.236, *P* = 0.0034; for CC1^+^, *t*_(4)_ = 16.92, *P* < 0.0001; for NG2^+^, *t*_(4)_ = 3.634, *P* = 0.0221. In B, for Olig2^+^, *t*_(10)_ = 5.08, *P* = 0.0005; for CC1^+^, *t*_(4)_ = 3.134, *P* = 0.0351; for NG2^+^, *t*_(17)_ = 6.387, *P* < 0.0001. In C, for Olig2^+^, *t*_(10)_ = 1.106, *P* = 0.295; for CC1^+^, *t*_(4)_ = 0.9848, *P* = 0.381; for NG2^+^, *t*_(9)_ = 1.062, *P* = 0.316. (**D**-**F**) Double immunostaining results of Olig2 and Ki67 in the corpus callosum of indicated mice at indicated ages. Arrows, representative double positive nuclei. (**G**) Statistic results of densities of proliferating OL lineage cells (Olig2^+^Ki67^+^) examined in indicated mice at indicated ages. Data were from immunostaining of 3-4 mice for each group, presented as mean ± s.e.m., and analyzed by two-tailed unpaired *t* test. For *Sox10*-ErbB2^V664E^ (SB) and control mice (Ctrl), *t*_(4)_ = 3.924, *P* = 0.0172. For *Sox10*-dnEGFR (SE) and Ctrl, at P35, *t*_(6)_ = 0.5042, *P* = 0.632; at P65, *t*_(6)_ = 2.963, *P* = 0.0252. For *Plp*-dnEGFR (PE) and Ctrl, at P35, *t*_(4)_ = 0.1136, *P* = 0.9151; at P65, *t*_(4)_ = 0.6191, *P* = 0.569. (**H** and **I**) Apoptotic cells (TUNEL^+^, white arrows) in the corpus callosum of *Sox10*-dnEGFR mice (H), or *Plp*-dnEGFR mice (I), were as few as that in littermate controls at indicated ages.

Neither differences in Olig2^+^, CC1^+^, NG2^+^, or Ki67^+^Olig2^+^ cell densities (Figure 5C,F,G; Figure 5-figure supplement 1C; Figure 5-figure supplement 2D-F), nor in TUNEL^+^ cells (Figure 5I), were observed in white matter of *Plp*-dnEGFR mice and littermate controls at P35 or P65. Different cellular and histological phenotypes in *Sox10*-dnEGFR and *Plp*-dnEGFR mice consolidated again the different targeting specificities of *Sox10^+/^*^rtTA^ and *Plp*-tTA. Moreover, the conflicting observations that the numbers of post-mitotic OLs (CC1^+^) increased but myelin thickness reduced in the brain of *Sox10*-dnEGFR mice (Figure 4E and Figure 5B), suggested that ErbB inhibition in OPC-NFOs had significantly impaired the myelinating capacity of OLs.

Given that *Sox10*-dnEGFR and *Sox1*0-ErbB2^V664E^ mice both exhibited hypomyelination, they should share a molecular or cellular deficit in myelination. We performed RNA-seq analyses of subcortical white matter tissues and identified 68 genes which exhibited similar expression tendencies in *Sox10*-ErbB2^V664E^ and *Sox10*-dnEGFR mice (Figure 6A). This group of genes have potential to regulate CNS myelination. Notably, in addition to *Gsn* and *Itgb4* that have been identified as characteristic genes for myelinating OLs (*Zhang et al., 2014*), *Enpp6*, *Itpr2*, and *Slc12a2* as characteristic genes for NFOs also exhibited significantly reduced expression in both mouse lines, supporting the notion that NFO deficiency contributes to hypomyelination. We examined the distribution of *Enpp6*-expressing cells by *in situ* hybridization (*Xiao et al., 2016*), and found that *Enpp6*^+^ cell numbers were indeed reduced in the corpus callosum of *Sox10*-dnEGFR mice at P35, although the reduction became indistinguishable for mice at P65 (Figure 6C,E). We further examined NFOs by immunostaining for TCF4, and found TCF4^+^ cell numbers were also reduced in the corpus callosum of *Sox10*-dnEGFR mice at P35, although the reduction became indistinguishable for mice at P65 (Figure 6G,I). Therefore, NFO differentiation was impaired shortly after ErbB inhibition, although it took 44 days to result in obvious hypomyelination in *Sox10*-dnEGFR mice. TCF4^+^ and *Enpp6*^+^ cell numbers were also reduced in *Sox10*-ErbB2^V664E^ mice (Figure 6B,F,E,I), which were due to the shortage of OPCs for differentiation.

**Figure 6.**
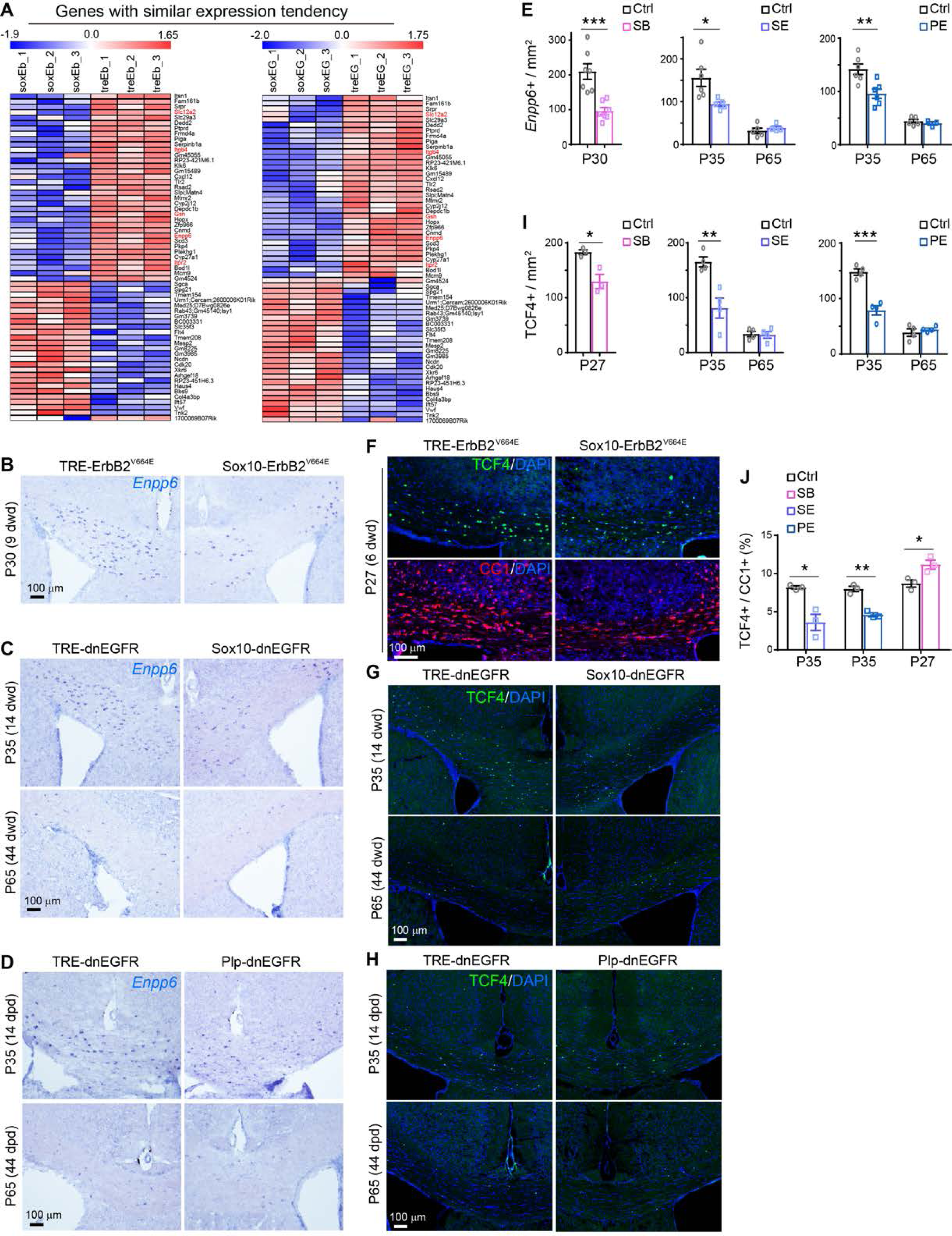
ErbB activation positively regulates NFO differentiation. (**A**) RNA-seq was performed for white matter tissues isolated from *Sox10*-ErbB2^V664E^ mice at P30 with 9 dwd, or *Sox10*-dnEGFR mice at P35 with 14 dwd, and their littermate controls. For each mouse group we sequenced three pairs of samples and identified 2298 genes that had altered expression in the white matter of *Sox10*-ErbB2^V664E^ mice (Figure 6-Source data 1), as well as 1184 genes in that of *Sox10*-dnEGFR mice (Figure 6-Source data 2). By comparing the two groups of genes, 68 genes with similar expression tendencies in the white matter of *Sox10*-ErbB2^V664E^ (soxEb *vs* treEb) and *Sox10*-dnEGFR (soxEG *vs* treEG) mice were identified. Heat maps of Z value of each gene are presented. The raw RNA-seq data have been deposited in the GEO and SRA database and can be found at GEO: GSE123491. (**B**-**D**) *In situ* hybridization results of *Enpp6* in the corpus callosum of *Sox10*-ErbB2^V664E^ mice (B), *Sox10*-dnEGFR mice (C), or *Plp-*dnEGFR mice (D), and their littermate controls at indicated ages. (**E**) Statistic results of *Enpp6*^+^ cell densities examined in indicated mice at indicated ages. Data were from repeated experiments of 3 mice for each group, presented as mean ± s.e.m., and analyzed by two-tailed unpaired *t* test. For *Sox10*-ErbB2^V664E^ mice (SB) and control mice (Ctrl) at P30, *t*_(12)_ = 4.638, *P* = 0.0006. For *Sox10*-dnEGFR mice (SE) and Ctrl, at P35, *t*_(9)_ = 2.704, *P* = 0.024; at P65, *t*_(7)_ = 0.8322, *P* = 0.433. For *Plp*-dnEGFR mice (PE) and Ctrl, at P35, *t*_(10)_ = 3.45, *P* = 0.0062; at P65, *t*_(6)_ = 0.7724, *P* = 0.469. (**F**-**H**) Immunostaining results of TCF4 in the corpus callosum of *Sox10*-ErbB2^V664E^ mice (F), *Sox10*-dnEGFR (G), or *Plp*-dnEGFR (H), with their littermate controls at indicated ages. (**I**) Statistic results of TCF4^+^ cell densities examined in indicated mice at indicated ages. Data were from repeated experiments of 3 mice for each group, presented as mean ± s.e.m., and analyzed by two-tailed unpaired *t* test. For *Sox10*-ErbB2^V664E^ mice (SB) and control mice (Ctrl) at P27, *t*_(4)_ = 3.883, *P* = 0.0178. For *Sox10*-dnEGFR mice (SE) and Ctrl, at P35, *t*_(6)_ = 4.107, *P* = 0.006; at P65, *t*_(6)_ = 0.1948, *P* = 0.852. For *Plp*-dnEGFR mice (PE) and Ctrl, at P35, *t*_(6)_ = 6.776, *P* = 0.0005; at P65, *t*_(6)_ = 0.7845, *P* = 0.463. (**J**) Ratio of TCF4^+^ to CC1^+^ cell numbers in indicated mice at indicated ages. Data were from 3 mice for each group, presented as mean ± s.e.m., and analyzed by two-tailed unpaired *t* test. For *Sox10*-dnEGFR mice (SE) at P35, *t*_(4)_ = 4.251, *P* = 0.0131; for *Plp*-dnEGFR mice (PE) at P35, *t*_(4)_ = 7.762, *P* = 0.0015; for *Sox10*-ErbB2^V664E^ (SB) mice at P27, *t*_(4)_ = 3.322, *P* = 0.029.

Interestingly, we also observed lowered TCF4^+^ and *Enpp6*^+^ cell numbers in *Plp*-dnEGFR mice at P35 (Figure 6D,E,H,I). *Plp*-tTA targeted a fraction of NFOs (Figure 2E-I). The different myelination states and similar NFO number reduction in *Sox10*-dnEGFR and *Plp*-dnEGFR mice suggest that, besides regulating myelinating capability of NFOs, ErbB signaling separately regulates another aspect of NFO differentiation, *i.e*., the transition time from NFO to MO stage. Indeed, in contrast to that in *Sox10*-dnEGFR and *Plp*-dnEGFR mice, the ratio of TCF4^+^ to CC1^+^ cell densities increased in *Sox10*-ErbB2^V664E^ mice (Figure 6J), which may suggest a prolonged transition to the MO stage for NFOs with ErbB activation.

### ErbB inhibition in MOs disrupts cognitive function in the absence of myelin alteration

OLs also offer essential trophic support to neurons in addition to forming myelin (*Nave and Werner, 2014*). We compared the behavioral performance of *Sox10*-dnEGFR mice, with hypomyelination in most brain regions, and *Plp*-dnEGFR mice, with normal myelin, to investigate whether disrupting ErbB signaling in MOs induces deficits other than dysmyelination. *Sox10*-dnEGFR mice performed worse than control mice in the rotarod test (Figure 7A), and were slightly hypoactive in the open field test (Figure 7B). Nevertheless, they performed normally in the central/peripheral zone analysis for assessment of anxiety, stereotyped behavior analysis and social interaction analysis for potential autistic-like phenotype, prepulse inhibition analysis for sensory gating, as well as forced swim and tail suspension tests for depression (Figure 7-figure supplement 1A-E). Interestingly, *Plp*-dnEGFR mice performed normally, similar to the controls in most tests, except exhibiting a subtle hyperactivity in the open field test (Figure 7C,D and Figure 7-figure supplement 1F-J). The different results from the two mouse lines implied that the impaired motor coordination could be attributed to the hypomyelination in the CNS.

**Figure 7.**
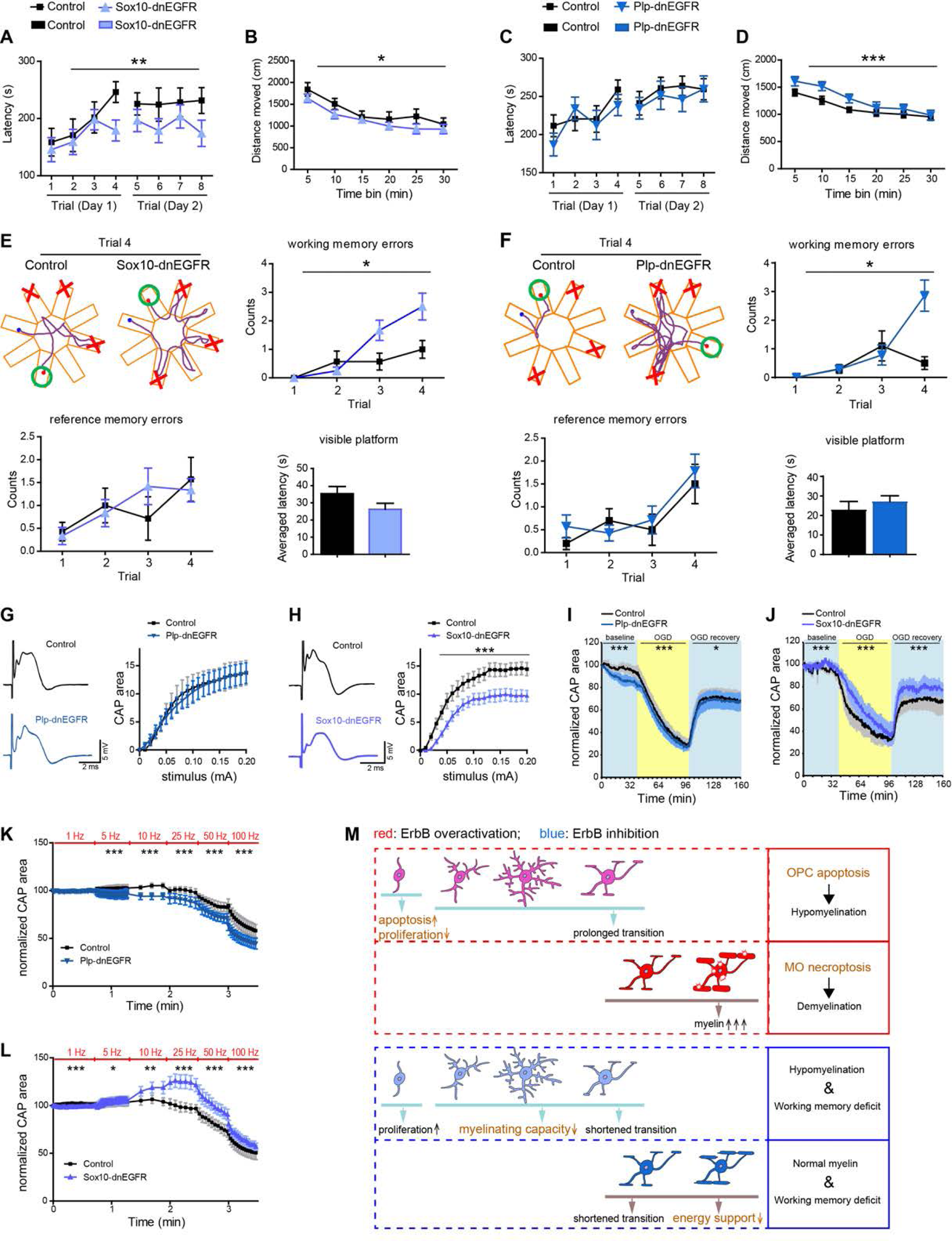
ErbB inhibition in MOs suppresses axonal conduction under energy stress and impairs working memory in the absence of myelin alteration. (**A** and **C**) Rotarod test, n = 12 mice for *Sox10*-dnEGFR and n = 12 mice for controls (two-way ANOVA test, *F*_(1, 176)_ = 7.824, *P* = 0.0057), while n = 12 mice for *Plp*-dnEGFR and n = 13 mice for controls (two-way ANOVA test, *F*_(1, 184)_ = 1.4, *P* = 0.238). (**B** and **D**) Open field tests, n = 13 mice for *Sox10*-dnEGFR and n = 11 mice for controls (two-way ANOVA test, *F*_(1, 132)_ = 6.302, *P* = 0.013), while n = 14 mice for *Plp*-dnEGFR and n = 19 mice for controls (two-way ANOVA test, *F*_(1, 186)_ = 12.17, *P* = 0.0006). (**E** and **F**) Eight-arm radial water maze test, n = 12 mice for *Sox10*-dnEGFR and n = 7 mice for controls (for working memory error, two-way ANOVA test, *F*_(1, 68)_ = 6.334, *P* = 0.014; for reference memory error, two-way ANOVA test, *F*_(1, 68)_ = 0.0423, *P* = 0.838; for visible platform, two-tailed unpaired *t* test, *t*_(17)_ = 1.137, *P* = 0.271), while n = 14 mice for *Plp*-dnEGFR and n = 10 mice for controls (for working memory error, two-way ANOVA test, *F*_(1, 88)_ = 4.782, *P* = 0.0314; for reference memory error, two-way ANOVA test, *F*_(1, 88)_ = 0.498, *P* = 0.482; for visible platform, two-tailed unpaired *t* test, *t*_(22)_ = 0.8254, *P* = 0.418). Data were presented as mean ± s.e.m.. In illustrative examples of the travel pathways of indicated mice, green circles indicate the last arms with a hidden platform, while red crosses indicate the arms with used platforms in the past three trials. (**G** and **H**) Axonal excitability is similar in *Plp*-dnEGFR optic nerves and control nerves (G), but decreased in *Sox10*-dnEGFR optic nerves in comparison with controls (H). CAPs of optic nerves generated by electrical stimuli with intensities at stepped increase (0-0.2 mA) were recorded *ex vivo*. Data were from 3-7 optic nerves of 3-5 mice for each group, presented as mean ± s.e.m., and analyzed by two-way ANOVA. In G, *F*_(1, 202)_ = 0.2118, *P* = 0.646. In H, *F*_(1, 139)_ = 102.1, *P* < 0.0001. Representative maximal CAPs for each group are shown at the left. (**I** and **J**) The CAP decline induced by OGD is slightly accelerated and aggravated in *Plp*-dnEGFR optic nerves (I), but decelerated and attenuated in *Sox10*-dnEGFR optic nerves (J). OGD was started for the recorded nerves after 1-hr baseline stimulation, and stopped after another hour by restoring the bathing media to oxygenated ACSF. Initial CAPs were recorded after 30-min baseline stimulation. The areas under CAPs were measured and normalized to the initial levels. Data were from 4-8 optic nerves of 3-5 mice for each group, presented as mean ± s.e.m., and analyzed by two-way ANOVA. In I, for baseline, *F*_(1, 1572)_ = 320.7, *P* < 0.0001; for OGD, *F*_(1, 2280)_ = 47.95, *P* < 0.0001; for recovery, *F*_(1, 2040)_ = 5.896, *P* = 0.015. In J, for baseline, *F*_(1, 732)_ = 87.06, *P* < 0.0001; for OGD, *F*_(1, 1140)_ = 196.6, *P* < 0.0001; for recovery, *F*_(1, 1020)_ = 173.1, *P* < 0.0001. (**K** and **L**) Neuronal activities with frequency at 5-100Hz increased the CAP decline in *Plp*-dnEGFR optic nerves (K), but increased the CAPs at 10-25Hz and slowed the CAP decline at 50-100Hz in *Sox10*-dnEGFR optic nerves (L), in comparison with their controls, respectively. Data were from 4-8 optic nerves of 3-5 mice for each group, presented as mean ± s.e.m., and analyzed by two-way ANOVA. In K, for 1Hz, *F*_(1, 270)_ = 0.076, *P* = 0.783; for 5Hz, *F*_(1, 1044)_ = 147.5, *P* < 0.0001; for 10Hz, *F*_(1, 27)_ = 17.64, *P* = 0.0003; for 25Hz, *F*_(1, 54)_ = 14.26, *P* = 0.0004; for 50Hz, *F*_(1, 90)_ = 19.82, *P* < 0.0001; for 100Hz, *F*_(1, 135)_ = 52.82, *P* < 0.0001. In L, for 1Hz, *F*_(1, 390)_ = 52.7, *P* < 0.0001; for 5Hz, *F*_(1, 1508)_ = 4.194, *P* = 0.041; for 10Hz, *F*_(1, 39)_ = 8.352, *P* = 0.0063; for 25Hz, *F*_(1, 78)_ = 50.46, *P* < 0.0001; for 50Hz, *F*_(1, 130)_ = 46.3, *P* < 0.0001; for 100Hz, *F*_(1, 195)_ = 19.59, *P* < 0.0001. (**M**) Schematic illustration of the pathophysiological consequences induced by ErbB receptor dysregulation in different OL stages as well as the pathogenic mechanisms (emphasized in brown).

We further tested these mice in the eight-arm radial water maze, a paradigm analyzing working memory capacity. It is known that myelin integrity is fundamental to cognitive performance of patients (*Kujala et al., 1997*). Moreover, although ErbB3/ErbB4 double knock-out does not induce myelin alteration in the CNS during early postnatal development (*Brinkmann et al., 2008*), a study of specifically depleting ErbB3 in mice from P19 has associated CNS hypomyelination with working memory deficits in adult mice (*Makinodan et al., 2012*). However, not only *Sox10*-dnEGFR mice, which had CNS hypomyelination, but also *Plp*-dnEGFR mice, which did not have myelin alteration, had significantly more working memory errors than control mice (Figure 7E,F and Figure 7-Video 1). Note that they had normal eyesight as performed in the visible platform test, as well as similar reference memory errors that indicated unaltered spatial recognition and memory (Figure 7E,F). This phenotype in *Plp*-dnEGFR mice reveals that working memory deficiency can be caused directly by ErbB inhibition in MOs through a myelination-independent mechanism.

### ErbB inhibition in MOs suppresses axonal conduction under energy stress

To determine what kind of function was impaired in white matter tracts of *Plp*-dnEGFR mice, we acutely isolated the optic nerves from adult mice (P90-P110) and recorded electrical stimulus-evoked compound action potentials (CAPs). The areas under CAPs, which are proportional to the total number of excited axons, indicate the nerve conduction. We found comparable areas under CAPs in *Plp*-dnEGFR optic nerves and control nerves responding to stimuli of the same intensity (Figure 7G). The maximal CAPs, which represent that all axons in the nerves are excited, were similar in *Plp*-dnEGFR optic nerves and control nerves (Figure 7G). In contrast, they were reduced in *Sox10*-dnEGFR optic nerves as compared with littermate controls (Figure 7H). These results reflected that the basic axonal conduction was not affected in *Plp*-dnEGFR white matter tracts, whereas it was impaired in *Sox10*-dnEGFR white matter tracts that exhibited hypomyelination.

In addition to myelin, macroglial metabolites are important for axonal conduction maintenance under conditions of energy deprivation (*Funfschilling et al., 2012; Saab et al., 2016; Trevisiol et al., 2017*). We challenged the optic nerves by incubating them in the oxygen-glucose deprivation (OGD) condition for 60 min. CAPs fell gradually in control optic nerves, and finally fell to 30% of the initial levels (Figure 7I,J). However, for *Plp*-dnEGFR optic nerves in the OGD condition, CAP failure was slightly accelerated and aggravated (Figure 7I). Contrarily, for *Sox10*-dnEGFR optic nerves under the same condition, CAP failure was decelerated and attenuated (Figure 7J). When the glucose and oxygen levels in the bathing medium were restored, CAPs in control optic nerves and *Plp*-dnEGFR optic nerves recovered to 60% of the initial levels (Figure 7I,J). However, in *Sox10*-dnEGFR optic nerves, CAPs recovered to 80% of the initial levels (Figure 7J).

It is notable that continuous electrical stimulation caused a baseline CAP decline in *Plp*-dnEGFR optic nerves, whereas a baseline CAP enhancement in *Sox10*-dnEGFR optic nerves, before the OGD (Figure 7I,J). Therefore, we further examined the axonal conduction under a physiological condition with increasing energy demands generated by neuronal activities (*Saab et al., 2016; Trevisiol et al., 2017*). By stimulating the control optic nerves with several trains of short bursts with frequency increased from 1 to 100 Hz, we confirmed that the low frequency stimulation (5-25Hz) has only minor influence on the CAPs, whereas the high frequency stimulation (50-100Hz) exhausts axonal energy and results in CAP decline (Figure 7K,L). For *Sox10*-dnEGFR optic nerves, intriguingly, the 5-25Hz electrical stimuli amplified CAPs and the 50-100Hz stimuli induced smaller CAP decay than that in control nerves (Figure 7L). In contrast, in *Plp*-dnEGFR optic nerves, either group of stimuli significantly aggravated the CAP decay (Figure 7K).

These results showed that *Sox10*-dnEGFR white matter tracts exhibited resistance to energy stress induced by both pathological (OGD) and physiological (neuronal activities) conditions. This may be ascribed to increased OL numbers, as that OLs are an essential venue for glycolysis and energy substrate production in support of axonal conduction (*Funfschilling et al., 2012*). In contrast, *Plp*-dnEGFR white matter was deficient in the maintenance of axonal conduction, especially under physiological energy stress. It is notable that MO numbers were not altered in *Plp*-dnEGFR mice that have ErbB inhibition in MOs, whereas ErbB receptors were not inhibited in MOs of *Sox10*-dnEGFR mice that have increased MOs (Figure 5B,C). Therefore, the opposite results of *Sox10*-dnEGFR and *Plp*-dnEGFR optic nerves in the energy challenging studies reveal that ErbB inhibition in MOs impairs the glia-axon energy coupling efficiency within electrically active neural circuits, which can compromise the cognitive function in *Plp*-dnEGFR mice in the absence of myelin alteration (Figure 7F).

## Discussion

Our results demonstrate that both ErbB3/ErbB4 receptors binding to the NRG family ligands and EGFR binding to the EGF family ligands are functional in adolescent and adult OLs. With the discovery of two valuable *in vivo* research mouse tools that differentially target OLs at MO and OPC-NFO stages, we reveal that NRG-ErbB and EGF-ErbB signaling cooperate in OPCs, NFOs, and MOs to simultaneously regulate myelination and axonal energy supporting functions. Aberrant ErbB activation or inhibition causes white matter abnormalities with distinct pathological characteristics and biological markers (Figure 7M).

ErbB overactivation is pathogenetic in MOs through inducing myelin overproduction and MO necroptosis, which results in demyelination followed by pathological changes including axon degeneration, OPC regeneration, astrogliosis and microgliosis. Notably, ErbB overactivation in OPCs induces apoptosis, without stimulating inflammatory pathological responses in the brain (Figures 1 and 3). Caspase-8 activation has been reported to be the key event to determine apoptotic fate of cells (*Oberst et al., 2011*), and defective activation of caspase-8 is critical for RIP1/RIP3/MLKL signaling to induce OL necroptosis in multiple sclerosis (*Ofengeim et al., 2015*). A cell-type specific RNA-sequencing transcriptome analysis suggests that caspase-8 is minimally expressed in post-mitotic OLs but is detectable in OPCs (*Zhang et al., 2014*), which may determine MO necroptosis but OPC apoptosis under continuous ErbB activation. Interestingly, studies on genetically modified mice that overexpressing hEGFR in OL lineage cells, or overexpressing NRG1 Type I or Type III in neurons, did not report myelin pathogenesis (*Aguirre et al., 2007; Brinkmann et al., 2008*). Nevertheless, mice with overactivation of the ErbB downstream signaling in OL lineage cells exhibit myelin and axonal pathology (*Harrington et al., 2010; Ishii et al., 2016*). *Olig2*-cre;Pten^flox/flox^ mice that overactivate PI3K/Akt signaling in OL lineage cells have loosened myelin lamellae in the spinal cord at 14 weeks and axonal degeneration in the cervical spinal cord fasciculus gracilis at 62 weeks (*Harrington et al., 2010*). *Plp*-CreER;Mek/Mek mice, which overexpress a constitutively activated MEK, a MAPK kinase, in OL lineage cells with tamoxifen induction, show demyelination in the spinal cord 3 months after MAPK (Erk) overactivation is induced (*Ishii et al., 2016*). Devastating effects of ErbB2^V664E^ in OLs may be due to its potent promotion of endogenous ErbB activation and multiple downstream signaling. Nevertheless, observations in the present study and in *Olig2*-cre;Pten^flox/flox^ and *Plp*-CreER;Mek/Mek mice corroborate the concept that constitutively activating ErbB signaling in OL lineage cells is pathogenetic, even though it may take a long time for moderate activation to result in pathological symptoms.

The profound demyelination or hypomyelination in *Plp*-ErbB2^V664E^ and *Sox10*-ErbB2^V664E^ mice should have disrupted many brain functions, although we could not evaluate these functions in the two strains by behavioral tests due to their severe motor dysfunction (Figure 1B,M). Intriguingly, a battery of behavioral tests for *Sox10*-dnEGFR and *Plp*-dnEGFR mice only revealed working memory deficits for both of them, except for the impaired motor coordination in *Sox10*-dnEGFR mice that have moderate hypomyelination (Figure 7A-F and Figure 7-figure supplement 1A-J). The further analyses emphasize that, although endogenous ErbB activation is required for both NFOs and MOs, it is used for the control of myelination and glia-axon energy coupling, respectively. Thus, ErbB inhibition in OLs impairs cognitive functions *via* myelination-dependent and -independent mechanisms. *Plp*-dnEGFR mice are a good model to affirm the myelination-independent contributions of OLs to higher brain function. As exhibited in *Plp*-dnEGFR mice, dual inhibition of NRG-ErbB and EGF-ErbB signaling in MOs does not affect myelin or OL numbers in the adolescent and adult brains. However, endogenous ErbB activities in MOs are indispensable for the maintenance of axonal conduction under physiological energy stress (Figure 7I,K), which are important for neuronal circuit efficiency as well as cognitive performance (Figure 7F). It is interesting that axonal conduction under energy stress was enhanced in *Sox10*-dnEGFR white matter tracts (Figure 7J,L), although it remains unclear whether the improved energy supplementation alleviates or aggravates the cognitive deficits in *Sox10*-dnEGFR mice (Figure 7E). Multiple questions remain unanswered, such as ErbB signaling regulates glucose metabolism in MOs or the transportation of energy metabolites from MOs to axons. ErbB dysregulation disrupts glutamatergic synaptic transmission in neurons (*Luo et al., 2014; Ting et al., 2011; Woo et al., 2007*). Glutamatergic synaptic transmission onto OLs was recently discovered to be essential for their energy substrate supply to axons (*Saab et al., 2016*), as well as for OL development (*Kougioumtzidou et al., 2017*). Therefore, it would be worth pursuing whether ErbB signaling regulates OL development or the trophic support from MOs to neurons by modulating glutamatergic synaptic transmission on OLs.

The roles of ErbB signaling in CNS myelination have long been debated because of the contradictory findings reported by different research groups. For example, ErbB3 has been reported to be dispensable for OL development (*Schmucker et al., 2003*), and ErbB3/ErbB4 double knockout does not result in CNS myelin alteration (*Brinkmann et al., 2008*). However, there are other reports that indicate inducing ErbB3 depletion by *Plp*-CreER in OL lineage cells from P19, not P36, results in adult hypomyelination (*Makinodan et al., 2012*). It is notable that ErbB3 has peaked expression during P15-P30 (Figure 1-figure supplement 1), and ErbB receptor manipulation in OPC-NFOs alters ErbB3/4 activities in white matter at P30-35 but not at P65 (Figure 1N and Figure 4B). Note that *Plp*-CreER is a mouse tool that can target OPCs and their progeny (*Guo et al., 2009*). Therefore, the hypomyelination in *Plp*-CreER;ErbB3^flox/flox^ mice is in line with our findings in *Sox10*-dnEGFR mice and reflects the positive role of ErbB signaling in NFO myelination during late postnatal development. EGFR is expressed stably in white matter during P20-P40 (Figure 1-figure supplement 1). The phosphorylation of EGFR is altered in white matter in all four mouse strains, which have ErbB receptor manipulation either in MOs or in OPC-NFOs (Figure 1C,N and Figure 4B,G). This suggests the general involvement of EGFR in OL function and development. The role of EGFR in CNS myelin development is supported by the report that transgenic mice with overexpression of hEGFR in all OL lineage cells (*CNP*-hEGFR) have enhanced myelin maturation, and hypomorphic EGFR mice (wa2) have delayed myelin maturation (*Aguirre et al., 2007*). *CNP*-hEGFR mice exhibit enhanced oligodendrogenesis in the subventricular zone, reflecting the function of EGFR in promoting neural progenitors to differentiate into OPCs during the early development (*Aguirre et al., 2007*). This is different from the pathological OPC regeneration revealed in the corpus callosum of *Plp*-ErbB2^V664E^ mice. It is notable that *CNP*-hEGFR increases the numbers of myelinated axons but not myelin thickness (*Aguirre et al., 2007*), which is different from the hypermyelination phenotype revealed in the CNS of NRG1-overexpressing mice (*Brinkmann et al., 2008*), suggesting that EGFR unlikely participates in myelin overproduction in MOs. It will be interesting to know whether the active EGFR, as revealed in *Plp*-dnEGFR mice (Figure 4G), is required for the trophic supportive function of MOs.

The negative regulation of ErbB activation on OPC proliferation and survival is unexpected because many *in vitro* studies have suggested otherwise. However, there is an interesting observation in transgenic mice *CNP*-dnErbB4, which are designed to overexpress a dominant negative ErbB4 mutant that specifically blocks the activities of ErbB3 and ErbB4 in all OL lineage cells. In this strain, post-mitotic OL numbers increase 40% in the corpus callosum although axons are hypomyelinated (*Roy et al., 2007*). Moreover, *Olig2*-cre;Pten^flox/flox^ mice, which have activation of the PI3K/Akt pathway in all OL lineage cells, exhibit hypermyelination but decreased OL densities in the developing corpus callosum (*Harrington et al., 2010*). These previously enigmagic observations are now well-explained by our findings that ErbB signaling plays different roles in OPCs and NFOs.

The white matter abnormalities observed in our mouse models are reminiscent of diverse myelin-related clinical and pathological characteristics in schizophrenic brains, including reduced white matter volume, decreased OL densities, reduced myelin gene products, apoptotic OLs, and damaged myelin (*Douaud et al., 2007; Fields, 2008; Hoistad et al., 2009; Uranova et al., 2011; Uranova et al., 2007*). Elevated ErbB activation has been repeatedly implicated in schizophrenia, and the increase could be caused by genetic factors (*Harrison and Law, 2006; Law et al., 2012*). To our knowledge, we are the first to reveal that ErbB overactivation can primarily induce oligodendropathy and myelin pathogenesis in white matter, providing a possible predisposition of a genetic variability in ErbB receptors or ligands to the white matter lesion. Notably, SNP8NRG243177 with T-allele, which increases NRG1 Type IV production (*Law et al., 2006*), is associated with the reduced white matter integrity (*McIntosh et al., 2008*) as well as increased psychotic symptoms (*Hall et al., 2006*) in schizophrenic patients.

Further, ErbB receptors and their ligands have been reported to reduce expression or lose function in some schizophrenic patients (*Harrison and Law, 2006; Mei and Nave, 2014*). Specific working memory deficits in *Sox10*-dnEGFR and *Plp*-dnEGFR mice firmly support that oligodendropathy can be a primary cause for the cognitive symptoms of schizophrenia. Moreover, myelin is very sensitive to environmental insults. Modest disruption of ErbB signaling by genetic mutation or SNPs is able to render myelin vulnerable to such insults, aggravating focal loss of connections under conditions of stress, ischemia, sleeplessness, trauma, etc. Therefore, OL dysfunction in patients, which is difficult to measure with current techniques, may eventually evolve into a detectable structural alteration in white matter that could contribute to another type of brain dysfunction. Collectively, our study provides novel insights into the pathophysiology of diseases initiated or aggravated within white matter.

## Materials and methods

### Animals

*Plp*-tTA transgenic mice (*Inamura et al., 2012*) were from the RIKEN BioResource Center (Stock No. RBRC05446). *Sox10^+/^*^rtTA^ mice were from Dr. Michael Wegner (University Erlangen-Nurnberg, Germany). Transgenic mice *TRE*-ErbB2^V664E^ (Stock No. 010577) and *TRE*-dnEGFR (Stock No. 010575) were from the Jackson Laboratory (*Chen et al., 2017*). Among ErbB1-4 receptors, ErbB2 that does not bind to any known ligand is the preferred partner to other ligand-bound ErbB members. ErbB2^V664E^ contains an amino acid mutation (Vla_664_/Glu_664_) within the transmembrane domain facilitating its dimerization with other ErbB receptors and potentiating their downstream signaling (*Chen et al., 2017*). DnEGFR, a dominant negative mutant of EGFR, is a truncated form of EGFR, losing the intracellular kinase domain but retaining the ability to form dimers with other ligand-bound ErbB members. When overexpressed, dnEGFR efficiently blocks the activation of any endogenous ErbB receptors under either NRG or EGF stimulation (*Chen et al., 2017*). Unless indicated, mice were housed under SPF conditions before experiments, in a room with a 12-hr light/dark cycle with access to food and water *ad libitum*. For biochemical and histological experiments, *Plp*-tTA;*TRE*-dnEGFR (*Plp*-dnEGFR), *Plp*-tTA;*TRE*-ErbB2^V664E^ (*Plp*-ErbB2^V664E^), *Sox10^+/^*^rtTA^;*TRE*-dnEGFR (*Sox10*-dnEGFR), or *Sox10^+/^*^rtTA^;*TRE*-ErbB2^V664E^ (*Sox10*-ErbB2^V664E^) mice with either sex and their littermate control mice with matched sex were used. For indicated behavioral tests, only male mice were used, while both male and female mice were used for the other behavioral tests because the results were not affected by sex difference. Animal experiments were approved by the Institutional Animal Care and Use Committee of the Hangzhou Normal University. For genotyping, the following primers were used: for *Plp*-tTA (630bp), PLPU-604 5’-TTT CCC ATG GTC TCC CTT GAG CTT, mtTA24L 5’-CGG AGT TGA TCA CCT TGG ACT TGT; for Sox10*^+/^*^rtTA^ (618bp), sox10-rtTA1 5’-CTA GGC TGT CAG AGC AGA CGA, sox10-rtTA2 5’-CTC CAC CTC TGA TAG GT CTT G; for *TRE*-dnEGFR (318bp), 9013 5’-TGC CTT GGC AGA CTT TCT TT, 7554 5’-ATC CAC GCT GTT TTG ACC TC; for *TRE*-ErbB2^V664E^ (625bp), 9707 5’-AGC AGA GCT CGT TTA GTG, 9708 5’-GGA GGC GGC GAC ATT GTC.

### Tet-Off or Tet-On treatment of mice

Mice with Tet-system contain genes of tetracycline-controlled transcriptional activator (tTA) or reverse tetracycline-controlled transcriptional activator (rtTA) driven by cell-specific promoters. When fed with Dox, these mice are able to switch on or off expression of a gene under the control of *tetracycline-responsive element* (*TRE*), specifically in rtTA- or tTA-expressing cells, which are called ‘Tet-on’ or ‘Tet-off’, respectively. The offspring of *Sox10*^+/rtTA^ during the indicated periods were fed with Dox (2 mg/mL × 10 mL/day from P21 to P35, and 1 mg/mL × 10 mL/day from P35 to indicated test day) in drinking water to induce the expression of ErbB2^V664E^ or dnEGFR in *Sox10*-ErbB2^V664E^ and *Sox10*-dnEGFR mice, respectively (Tet-On). For the offspring of *Plp*-tTA, Dox was given (Tet-off) from the embryonic day (through pregnant mothers) to their weaning day at P21 to inhibit the expression of ErbB2^V664E^ or dnEGFR during this period in *Plp*-ErbB2^V664E^ or *Plp*-dnEGFR mice (0.5 mg/mL × 10 mL/day of Dox before P21). Water bottles were wrapped with foil to protect Dox from light. All used littermate control mice were treated the same.

### Stereotactic injection of AAV viruses

pAAV-*TRE*-EYFP plasmids (Addgene) were packaged as AAV2/9 viruses, and produced with titers of 2.0E+13 particles per mL by OBio (Shanghai, China). Mice were anesthetized by 1% pentobarbital (50 mg/kg, i.p.) and mounted at stereotaxic apparatus (RWD68025). AAV-*TRE*-EYFP (2 μL) was injected into the corpus callosum (from bregma in mm, M-L: ±1.2, A-P: +0.5, D-V: -2.2) under the control of micropump (KDS310) at a speed of 0.05 μL/min. Injecting needles (Hamilton NDL ga33/30 mm/pst4) were withdrawn 10 min after injection. Infected brains were isolated 1 or 2 days later and brain slices were immunostained with anti-GFP antibody to enhance the visualization of the reporter protein.

### Electron Microscopy

Mice were anesthetized and transcardially perfused with 4% sucrose, 4% paraformaldehyde (PFA) and 2% glutaraldehyde in 0.1 M phosphate buffer (PB, pH 7.4). The brains, optic nerves, or sciatic nerves were isolated carefully. The corpora callosa and prefrontal cortices were further dissected carefully under stereoscope. Tissues were post-fixed overnight at 4°C in 1% glutaraldehyde in 0.1 M PB. Samples were washed by 0.1 M PB 24 hr later, and osmicated with 2% osmium tetroxide 30-60 min at 4°C, washed by 0.1 M PB and by deionized H_2_O at 4°C, and dehydrated in graded (50-100%) ethanol. Samples were incubated with propylene oxide and embedded with embedding resins. Ultrathin sections were stained with 2% uranyl acetate at 4°C for 30 min, and then photographed with Tecnai 10 (FEI). EM images were analyzed by Image J (NIH). To eliminate the bias on circularity, *g*-ratio of each axon was calculated by the perimeter of axons (inner) divided by the perimeter of corresponding fibers (outer). Axonal diameters were normalized by perimeters through equation: diameter = perimeter/π. This procedure allows for inclusion of irregularly shaped axons and fibers and helps to eliminate biased measurement of diameters based on circularity. For quantitative analysis, cross sections of each neural tissue were divided into 5 areas, and more than two images, randomly selected from each area, were examined.

### Immunofluorescence staining

Deeply anesthetized mice were transcardially perfused with 0.01 M PBS and then 4% PFA in 0.01 M PBS. Mouse brains were isolated and post-fixed in 4% PFA in 0.01 M PBS overnight at 4 °C, and then transferred into 20% and subsequently 30% sucrose in PBS overnight at 4 °C. Brains were then embedded in OCT (Thermo Fisher scientific) and sectioned into 20 μm on a cryostat sectioning machine (Thermo Fisher scientific, Microm HM525). Brain slices were incubated with blocking buffer (10% fetal bovine serum and 0.2% Triton-X-100 in 0.01 M PBS) for 1 hr at room temperature, and then incubated at 4 °C overnight with primary antibodies diluted in blocking buffer. The primary antibodies used were: GFP (1:500, Abcam, ab13970), CC1 (1:500, Abcam, ab16794), NG2 (1:200, Abcam, ab50009), Ki67 (1:400, Cell Signaling Technology, 9129), GFAP (1:2000, Millipore, MAB360), Iba1 (1:1000, Millipore, MABN92), TCF4 (1:500, Millipore, 05-511), Olig2 (1:500, Millipore, AB9610), TUJ1 (1:500, Sigma, T5076), RIP3 (1:500, QED, 2283), MLKL (1:500, Abgent, AP14272B). After washing three times with 0.1% Triton-X-100 in 0.01 M PBS, samples were incubated at room temperature for 1 hr with Alexa-488 or -594 secondary antibody, and then washed and mounted on adhesion microscope slides (CITOTEST) with fluorescent mounting medium. Nuclear labeling was completed by incubating slices with DAPI (0.1 μg/mL, Roche) at room temperature for 5 min after incubation with secondary antibodies. Except for the antibody against NG2, antigen retrieval in 0.01 M sodium citrate buffer (pH 6.0) at 80-90 °C for 10 min was necessary before primary antibody incubation for brain slices to achieve definitive signals. Images were taken by a Zeiss LSM710 confocal microscope or a Nikon Eclipse 90i microscope. For cell counting based on immunostaining results, soma-shaped immunoreactive signals associated with a nucleus was counted as a cell. The immunostaining intensity was measured by Image J with background subtraction.

### Luxol fast blue (LFB) staining

After sufficient washing with 0.01 M PBS, PFA-fixed brain slices were transferred into a mixture of trichloromethane and ethanol (volume ratio 1:1) for 10 min and then 95% ethanol for 10 min. They were next incubated in 0.2% Luxol fast blue staining solution (0.2 g Solvent blue 38, 0.5 mL acetic acid, 95% ethanol to 100 mL) at 60 °C overnight. In the next day, tissues were incubated for 5 min each in turn in 95% ethanol, 70% ethanol and ddH_2_O for rehydration, followed by incubation alternatively in 0.05% Li_2_CO_3_, 70% ethanol and ddH_2_O for differentiation until the contrast between the gray matter and white matter became obvious. After that, tissues were incubated for 10 min each in 95% and 100% ethanol to dehydrate, and then 5 min in dimethylbenzene to clear, before quickly mounting with neutral balsam mounting medium (CWBIO). All steps were operated in a ventilation cabinet. The LFB intensity in the corpus callosum was measured by Image J with background subtraction, and normalized to that of controls.

### TUNEL assay

Apoptotic cells were examined with terminal deoxynucleotidyl transferase (TdT)-mediated deoxyuridine triphosphate (dUTP) nick-end labeling (TUNEL) assay according to the manufacturer’s instructions (Vazyme; Yeasen). In brief, PFA-fixed brain slices were digested for 10 min by proteinase K (20 μg/mL) at room temperature. After washing twice with PBS, brain slices were incubated with Equilibration Buffer for 30 min at room temperature, and subsequently with Alexa Fluor 488-12-dUTP Labeling Mix for 60 min at 37°C. After washing with PBS three times, brain slices were stained with DAPI before being mounted under coverslips. For co-labeling of apoptotic nuclei in slices with immunofluorescence staining, TUNEL assay was performed after washing of the secondary antibody.

### Western blotting

Subcortical white matter tissues were isolated and homogenized. Homogenates in lysis buffer (10 mM Tris-Cl, pH 7.4, 1% NP-40, 0.5% Triton-X 100, 0.2% sodium deoxycholate, 150 mM NaCl, 20% glycerol, protease inhibitor cocktail) at a ratio of 2 mL per 100 mg tissue were lysed overnight in 4°C. Lysates were centrifuged at 12,000 g and 4°C for 15 min to get rid of the unsolved debris. Concentration of the supernatant was measured by BCA assay. Proteins in samples were separated by 6-12% SDS-PAGE, transferred to a Immobilon-P Transfer Membrane (Millipore), and then incubated with indicated primary antibodies diluted in blocking buffer at 4°C overnight after blocking by 5% non-fat milk solution in TBST (50 mM Tris, pH 7.4, 150 mM NaCl, 0.1% Tween 20) for 1 hr at room temperature. The primary antibodies used were: pErbB3 (1:2500, Abcam, ab133459), pErbB4 (1:2500, Abcam, ab109273), pErbB2 (1:2500, Abgent, AP3781q), EGFR (1:5000, Epitomics, 1902-1), pEGFR (1:2500, Epitomics, 1727-1), GAPDH (1:5000, Huabio, EM1101), MBP (1:1000, Millipore, MAB382), Olig2 (1:1000, Millipore, MABN50), ErbB3 (1:200, Santa Cruz Biotechnology, sc-285), ErbB4 (1:200, Santa Cruz Biotechnology, sc-283), ErbB2 (1:200, Santa Cruz Biotechnology, sc-284), RIP3 (1:2000, QED, 2283), MLKL (1:2000, Abgent, AP14272B). For antibodies against phosphorylated proteins, 10% fetal bovine serum was used as blocking buffer. Next day, the membranes were washed by TBST for three times and incubated with the secondary antibodies for 1 hr at room temperature. Membranes were washed again and incubated with Immobilon

Western Chemiluminescent HRPSubstrate (Millipore) for visualization of chemiluminescence by exposure to X-ray films or Bio-Rad GelDOCXR^+^ Imaging System. Intensities of protein bands were measured by Image J, and statistical analysis was performed after subtraction of the background intensity and normalization with controls in each batch of experiments.

### *In situ* hybridization

RNA *in situ* hybridization was performed as previously described (*Schaeren-Wiemers and Gerfin-Moser, 1993*) with minor modifications. Briefly, the 14 μm PFA-fixed brain sections were post-fixed in 4% PFA in PBS for 20 min, incubated in 2 μg/mL Proteinase K in 50 mM Tris-Cl (pH 7.4) with 5 mM EDTA at room temperature for 10 min, re-fixed in 4% PFA in PBS for another 10 min, and then acetylated in 1.33% triethanolamine and 0.25% acetic anhydride solution at room temperature for 10 min. The acetylated sections were washed and incubated in hybridization buffer (50% formamide, 0.25 mg/mL yeast RNA, 0.5 mg/mL herring sperm DNA, 5x Denhard’s, 5x SSC, Invitrogen) at room temperature for 1 hr, and then hybridized with 0.5 ng/μL digoxigenin-labeled *Enpp6* riboprobe in hybridization buffer at 65°C for 16 hr. The hybridized sections were washed three times in 0.2x SSC at 65°C for total 1 hr, and then were blocked with 10% sheep serum (Sigma-Aldrich) in solution I containing 100 mM pH 7.5 Tris-Cl with 0.15 M NaCl at room temperature for 1 hr, followed by incubation with alkaline phosphatase-conjugated anti-digoxigenin antibody (Roche) in the same solution at 4°C overnight. After washing three times with solution I for total 1 hr, and twice with developing buffer containing 100 mM pH 9.5 Tris-Cl, 0.1 M NaCl, 50 mM MgCl_2_ and 0.1% Tween-20, the sections were incubated with 2% NBT/BCIP solution (Roche) in the developing buffer at room temperature in the dark. The reaction was stopped by immersing the sections in PBS with 5 mM EDTA when appropriate signals were detected. To obtain mouse *Enpp6* riboprobes, a 1.3 kb fragment corresponding to Enpp6 mRNA (1400–2700 nt of NM_177304.4) was cloned into pBluescript II KS(-). The linearized plasmids were used as templates for *in vitro* transcription with T3 RNA polymerase (Promega) according to the manufacturer’s instructions.

### Real-time reverse transcription-PCR (RT-PCR)

Total RNA was extracted from isolated mouse white matter using TRIzol following manufacturer’s protocol. cDNA was synthesized by using the 5x All-In-One RT MasterMix (abmGood). Real-time PCR was performed in four repeats for each sample by using BrightGreen 2x qPCR MasterMix (abmGood) with the Bio-Rad CFX96 real-time PCR system as previously described (*Chen et al., 2017*). Relative mRNA levels were analyzed by software Bio-Rad CFX Manager 1.5. Transcripts of targeted genes were normalized to those of mouse 18S rRNA gene in the same samples. Primers for 18S rRNA were 5’-CGG ACA CGG ACA GGA TTG ACA and 5’-CCA GAC AAA TCG CTC CAC CAA CTA with a 94 bp PCR product. Primers for mouse EGFR gene and transgene dnEGFR were 5’-TCC TGC CAG AAT GTG AGC AG and 5’-ACG AGC TCT CTC TCT TGA AG with a 500 bp PCR product.

### RNA-Seq Analyses

Subcortical white matter tissues isolated from *Sox10*-dnEGFR and littermate controls, or *Sox10*-ErbB2^V664E^ mice and littermate controls, were used (three pairs for each group) for global transcriptome analysis by LC-Bio Co (Hangzhou, China). The final transcriptome was generated by Histat and StringTie. StringTie was used to estimate the expression level for mRNAs by calculating FPKM (Fragments Per Kilobase of exon model per Million mapped reads). Differentially expressed genes were identified by comparing FPKM of the mRNA reads from three sample pairs between *Sox10*-dnEGFR, or *Sox10*-ErbB2^V664E^ mice, and their littermate controls, by paired Student *t* test *via* MeV (MultiExperiment Viewer). Gene lists with significant difference (*P* < 0.05) in expression between *Sox10*-dnEGFR and littermate controls, or between *Sox10*-ErbB2^V664E^ and littermate controls, were compared, and genes with similar expression tendencies in *Sox10*-dnEGFR and *Sox10*-ErbB2^V664E^ mice were identified. Z value of these genes was calculated according to their FPKM by an equation “Z sample-i = [(log2(Signal sample-i)-Mean (Log2(Signal) of all samples)][Standard deviation (Log2(Signal) of all samples)]” and plotted as heat map by MeV. Gene Ontology (GO) term enrichment was analyzed by PANTHER Overrepresentation Test (Released 20171205) through http://geneontology.org with the significance estimated by Fisher’s Exact Test (FDR, false discovery rate).

### Behavioral Tests

*Plp*-ErbB2^V664E^ mice at P35 and *Sox10*-ErbB2^V664E^ mice at P30 after indicated Dox treatment were used in grid walking tests for motor function analysis. Behavioral analyses for *Sox10*-dnEGFR and littermate controls with Dox-feeding from P21, or *Plp*-dnEGFR mice and littermate controls with Dox-withdrawal from P21, were carried out with 12- to 16-week-old animals by investigators unaware of their genotypes. Tested mutant mice had littermate control mice with same sex. For PPI, social interaction, eight-arm radial water maze, forced swim and tail suspension, all tested mice were male. Animals were tested at a sequence of open field, social interaction, rotarod, PPI, eight-arm radial water maze, and then forced swim and tail suspension, to minimize the influence of stress on their behavioral performance. There were 2-day gaps between tests.

### Open field and stereotyped behavior analysis

Animals were placed in a chamber (30 cm × 30 cm × 34.5 cm) and their movements were monitored and traced by a tracking software EthoVision XT 12 (Noldus, The Netherland). Locomotive activity was measured and summated at 5-min intervals over a 30-min period. Frequency and cumulative duration of stereotyped behaviors observed during 30-min traveling in the open field, including grooming, hopping, rearing supported, and sniffing, were determined by EthoVision XT 12 and statistically analyzed. Anxiety of the animals was assessed by the differences of time that they spent in the central zone and peripheral zone during the 30 min.

### Rotarod

To evaluate the sensorimotor coordination, mice were placed on an accelerating rotarod (Mouse Rota-Rod NG, Harikul Science, UB47650, Italy) and assessed for ability to maintain balance on the rotating bar that accelerated from 4 to 40 rpm over a 5-min period. Mice were tested for 4 trials in the first day with 30-min gap between trials, and were tested for another 4 trials 24 hr later. Latency before fall from the rod was recorded.

### Prepulse inhibition (PPI) test

These tests were conducted in a sound-attenuated chamber (Panlab, LE116, Spain). Mice were placed in a Plexiglas restrainer mounted on a grid floor, and their startle responses were captured by a movement sensor and analyzed by a software Startle v1.2. Before the test, mice were allowed to habituate to the chamber with a 60 dB background white noise for 5 min, and to 4 times of auditory-evoked startle stimulating pulse (10000 Hz at 120 dB, 20 ms) with random 5-30 sec intervals. In the PPI test, mice were subjected to startle pulse trials (120 dB, 20 ms) or prepulse/pulse trials (20 ms 10000 Hz at 75, 80, or 85 dB with 100 ms interval before a 20 ms 120dB startle stimulus) with random 5-30 sec intervals between trials. Different trial types were presented randomly with each trial type presented 9-12 times, and no two consecutive trials were identical. Max startle response within 300 ms after onset of startle stimulus was recorded. PPI (%) was calculated according to the equation: [100 - (startle amplitude on prepulse-pulse trials/startle amplitude on pulse alone trials) ×100].

### Social interaction

Mice were placed in a square chamber (50 cm × 50 cm × 50 cm) with two small transparent boxes at two opposite corners. The chamber was dark and mouse movement was monitored by infrared camera and traced by Anymaze software (Stoelting, Wood Dale, Illinois). After 5 min habituation in the chamber, mice were returned to their home cages. Mice were placed into the same chamber 2 hr later, with one of the boxes holding a stranger mouse that could be seen and smelled through multiple holes in the box. Social interaction ability of the tested mice was determined by their traveling distances in the quadrant with the stranger mouse as compared with the traveling distances in the quadrant with an empty box.

### Eight-arm radial water maze

According to the previous report with modification (*Penley et al., 2013*), animals were trained in eight-arm radial water maze for two weeks, with four trials each day to search a hidden platform in each trial for escaping from the water at 20-22 °C. Four hidden platforms were placed at the end of a same set of arms for all the training and tests, as illustrated in Figure 7-figure supplement 1K. After a trial that mice reached a hidden platform, mice were returned to their home cages with towel and warming pads. There was a 30-sec gap between trials, and the visited platform was removed before the next trial. Mice aborted swimming during training were discarded. Two weeks later, the trained mice were tested for their working memory capacities that were represented by avoiding arms with visited platforms in previous trials. In the last trial of the test day, the animals had highest working memory load for they had to avoid swimming into three arms with platforms removed. First and repeat entries into any arm that previously had a platform were counted as working memory errors, and first entries into any arm that never had a platform were counted as reference memory errors which represent deficits in spatial recognition or long term memory. The day after test day, all the animals were tested in a simple visible platform task with 5 trials in a round pool, and each trial contained a visible platform placed at a different position 0.5-1.0 meter away from tested mice. The latency of mice reached the visible platform in each trial was averaged to assess their eyesight.

### Forced swim and Tail suspension

In the forced swim test, mice were forced to swim for 15 min in a cylinder with diameter at 11 cm, water depth at 30 cm and temperature at 22-24 °C. One day later, mice were forced to swim again for 5 min in the same cylinder and mouse movement was recorded and analyzed by Anymaze software. The tail suspension test was carried out 2 days later after the forced swim test, in which mice were suspended by using adhesive tape applied to the tail and videotaped for 5 min. Mouse movement during the 5 min was traced and analyzed by Anymaze software. For both tests, immobile period was defined by 70% of mouse bodies were motionless and lasted for at least 1 sec. Summation of immobile periods for each mouse was taken into statistical analysis.

### Grid walking test

Mice were placed on an elevated metal grid panel with each grid cell 5 × 0.8 cm^2^, and their movements were videotaped. The velocity of mouse movement and percentage of foot-slip steps in total steps were calculated to assess locomotor function of mice. Scores of foot slips reflect precise stepping, coordination of the four limbs, and accurate paw placement, indicating ability of animals in sensorimotor coordination.

### Electrophysiology

Following anesthesia and decapitation, optic nerves were isolated from mice and superfused with oxygenated artificial cerebrospinal fluid (ACSF) containing (in mmol/L): 119 NaCl, 2.5 KCl, 2.5 CaCl2,1.3 MgCl2, 1.25 NaH2PO4, 26 NaHCO3, and 10 glucose; pH 7.4. Optic nerve CAP recording methods were adopted and modified from the previous reports (*Saab et al., 2016*). Briefly, two ends of the optic nerves were attached by suction electrodes, which backfilled with ACSF and connected to an IsoFlex (AMPI) for stimulation or a MultiClamp 700B (Molecular Device) for recording. The recorded signal was amplified 50 times, filtered at 30 kHz, and acquired at 20-30 kHz. Data were collected and analyzed by pClamp 10.3 software (Molecular Devices). The optic nerves were equilibrated for at least 30 min in the perfusion chamber in normal ACSF at room temperature before experiments. All experiments were performed at room temperature.

### Maximal CAP recording

For each recorded nerve, stimulus pulse (100 µs duration) strength was adjusted with a stepped increase and finally to evoke the maximal CAP. CAPs were elicited 5 times at every step of the stimulus strength. After reaching the maximal CAP, the stimulus was increased an additional 25% for supramaximal stimulation to ensure the activation of all axons in the nerve. Note the supramaximal stimulation did not further change the CAPs. The areas under CAPs were calculated to determine the nerve conduction.

### Oxygen-glucose deprivation (OGD) assay

The assay was performed as previously described with modification (*Saab et al., 2016; Trevisiol et al., 2017*). During experiments, CAPs were evoked by the supramaximal stimulus every 20 sec. After 60-min stimulation of the baseline CAP in normal condition, OGD was induced for the nerves by switching bathing solution from oxygenated ACSF (saturated with 95% O2/5% CO2) to glucose-free ACSF (replaced with equimolar sucrose to maintain osmolarity) that was saturated with 95%N2/5%CO2. After 60-min OGD, oxygenated ACSF was restored and CAPs were recorded for another hour. CAPs recorded after 30-min baseline stimulation was taken as the initial CAPs. The effects of OGD on the nerve conduction and recovery were determined by normalizing the areas of CAPs recorded during OGD or recovery sessions to that of initial CAPs.

### Neural activity dependence assay

The protocol was modified from published reports (*Saab et al., 2016; Trevisiol et al., 2017*). Before the experiments, CAPs were recorded every 30 sec to obtain baseline with the stimulus pulse strength set at the supramaximal levels. To evaluate the conduction of optic nerves under increasing energy demands, we gradually increased the stimulating frequency from 1 to 100 Hz. Each stimulating frequency was applied for 30-60 sec. For 1 and 5 Hz stimuli, CAPs were continuously recorded and CAP areas were measured for each CAP. For 10 to 100 Hz, nerves were stimulated by a train of 100 stimuli, and rest for 1 sec before the next train of stimuli. CAP areas were sampled for the last four stimuli of each train and averaged as one data point. For the statistical analysis, CAP areas were normalized to the initial levels.

### Statistical analysis

Statistical analyses other than for RNA-seq data (described separately above) were performed using Prism (Graphpad) and presented as mean ± s.e.m.. For western blotting and LFB staining results, statistical analyses were performed after subtraction of the background intensity and normalization with controls in each batch of experiments to minimize the influences of batch-to-batch variations. Two-tailed unpaired Student’s *t* test was used for analysis between two groups with one variable, one-way ANOVA test was used for analysis among three or more groups with one variable, and two-way ANOVA test was used to determine difference among groups with two variables. Statistical significance was set at \**P* < 0.05, \*\**P* < 0.01, \*\*\**P* < 0.001.

## Data Availability

All data generated or analyzed during this study are included in the manuscript and supporting files. Source data files have been provided for all manuscript figures. Source data have been provided online at datadryad.org (https://doi.org/10.5061/dryad.jq2bvq87c). The accession number for the RNA-Seq data presented in this article is GEO: GSE123491.

## Acknowledgements

We thank Wanwan He, Kaiwei Zhang, Youguang Yang, and Shasha Zhang in Hangzhou Normal University for the assistance in EM image analyses, and Dr. Wanhua Shen for the assistance in electrophysiological experiments. We also thank Dr. Woo-Ping Ge in UT Southwestern for comments on the manuscript.

This work was supported by grants from National Natural Science Foundation of China (31371075, 31671070, and 31871030 to YT).

## Competing interests

The authors declare no competing interests.

---

**Figure 1-figure supplement 1.**
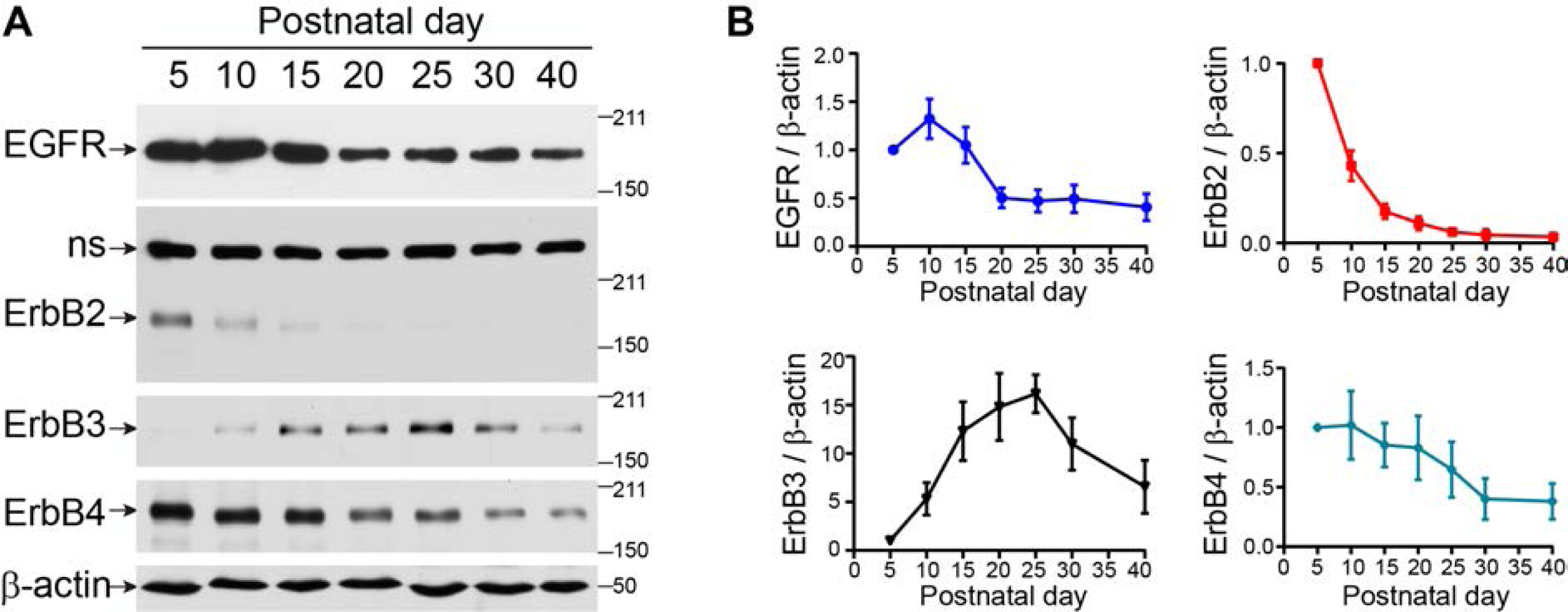
Expression of ErbB receptor members in white matter regions at different postnatal days. (**A**) Western blotting results of ErbB receptors in white matter regions isolated from wild type mice at different postnatal days. ns, non-specific band. (**B**) Quantitative data of the western blotting results were presented as mean ± s.e.m.. n = 3 independent experiments for each postnatal day. Data were normalized to P5 in each batch of experiments.

**Figure 1-figure supplement 2.**
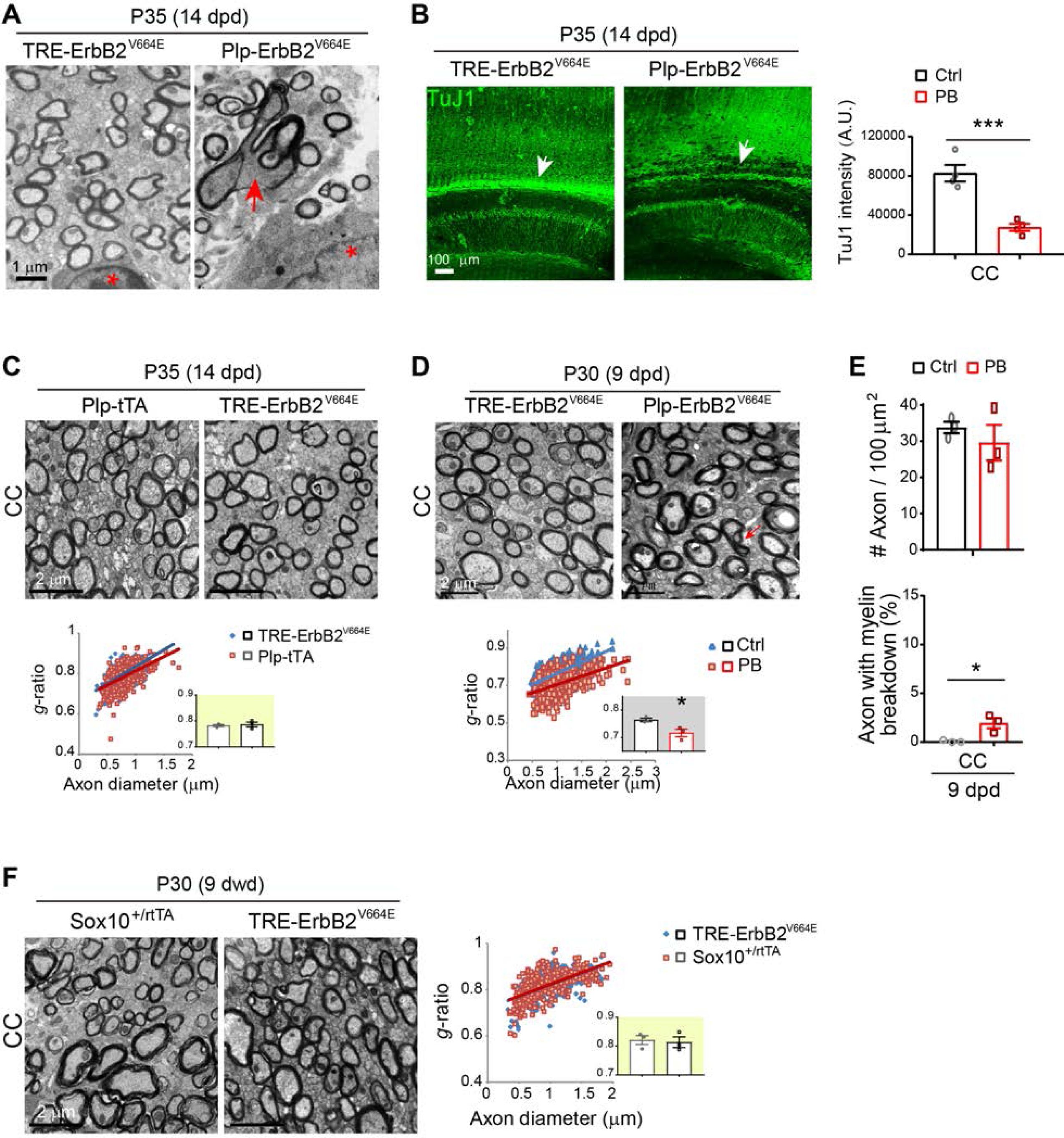
Unaltered myelin in the brains of *Plp*-tTA and *Sox10*^+/rtTA^ mice, but hypermyelination at 9 dpd and axonal pathology at 14 dpd in *Plp*-ErbB2^V664E^ mice. (**A**) Representative EM images showed that myelin sheath ruptured and broke down in *Plp*-ErbB2^V664E^ mice at 14 dpd (red arrow). Note the associated nuclei (red asterisk) showed no chromatin condensation and nucleation. (**B**) Dramatically reduced axons in the subcortical white matter of *Plp*-ErbB2^V664E^ mice at 14 dpd. Sagittal sections of *Plp*-ErbB2^V664E^ (PB) and littermate control mice (Ctrl) were immunostained by monoclonal antibody TuJ1. Quantitative data were presented as mean ± s.e.m., and analyzed by two-tailed unpaired *t* test. *t*_(6)_ = 6.019, *P* = 0.0009. (**C**) EM images of the corpus callosum (CC) of *Plp*-tTA and littermate *TRE*-ErbB2^V664E^ mice at P35 with 14 dpd. *g*-ratio was calculated for myelinated axons. Averaged *g*-ratio (inset) were presented as mean ± s.e.m., and analyzed by two-tailed unpaired *t* test. *t*_(4)_ = 0.4472, *P* = 0.678. (**D**) EM examination of axons in the midline of corpus callosum in *Plp*-ErbB2^V664E^ mice at 9 dpd. Note most axons remained intact despite that there were a few axons had myelin breakdown (red arrow). *g*-ratio was calculated for myelinated axons and averaged *g*-ratio for each mouse were analyzed by two-tailed unpaired *t* test (inset). *t*_(4)_ = 3.226, *P* = 0.0321. (**E**) The densities of myelinated axons as well as the percentages of axons with myelin breakdown in EM analysis of the midline of corpus callosum in *Plp*-ErbB2^V664E^ mice (PB) and littermate controls (Ctrl) at 9 dpd. Data were presented as mean ± s.e.m., and analyzed by two-tailed unpaired *t* test. For myelinated-axon density, *t*_(4)_ =0.805, *P* = 0.466. For axons with myelin breakdown, *t*_(4)_ = 3.567, *P* = 0.023. (**F**) EM images of the corpus callosum of *Sox10*^+/rtTA^ and littermate *TRE*-ErbB2^V664E^ mice at P30 with 9 dwd. *g*-ratio was calculated for myelinated axons. Averaged *g*-ratio (inset) were presented as mean ± s.e.m., and analyzed by two-tailed unpaired *t* test. *t*_(4)_ = 0.3042, *P* = 0.776.

**Figure 2-figure supplement 1.**
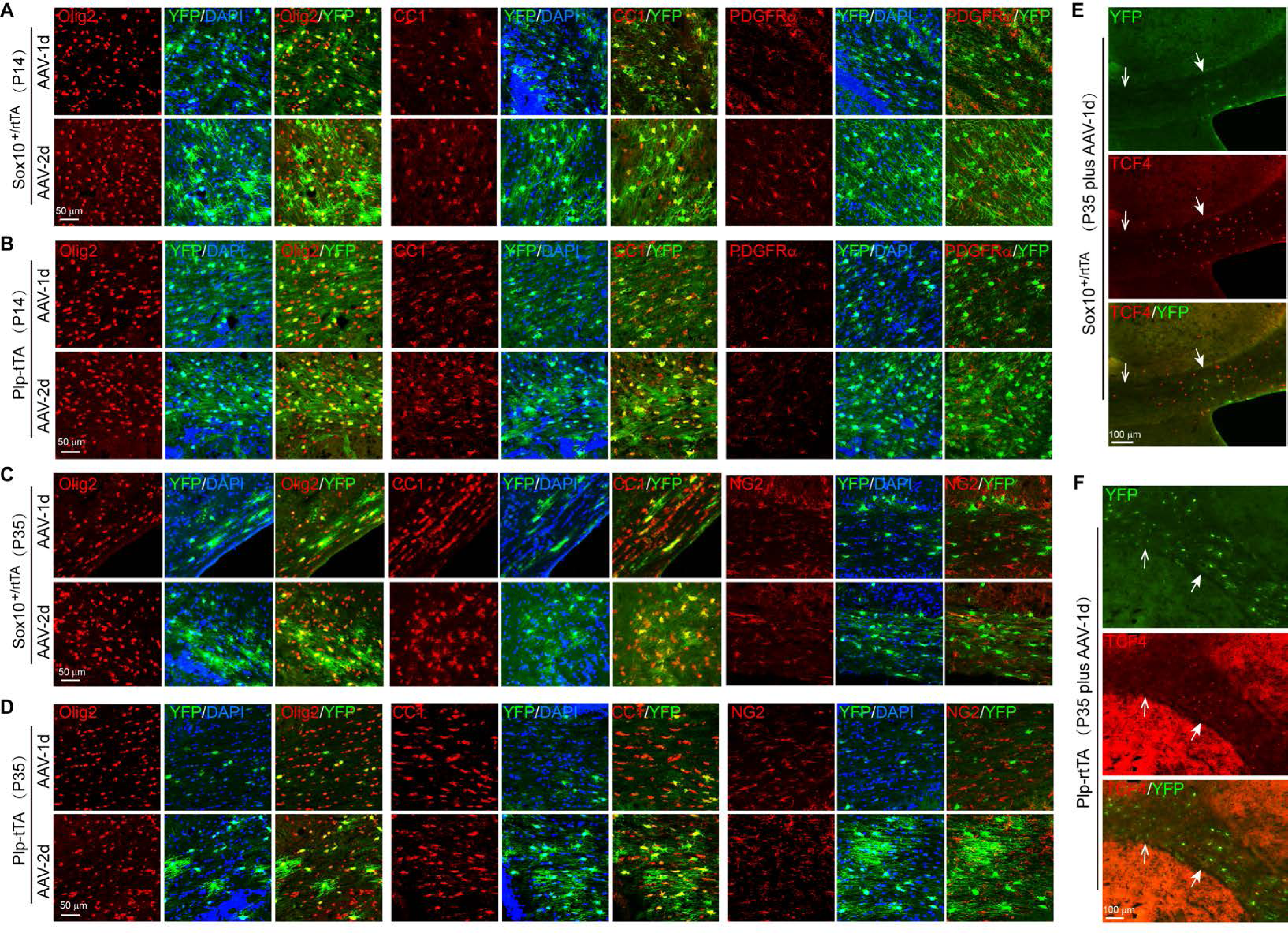
Targeting preferences on OPCs or post-mitotic OLs of *Plp*-tTA and *Sox10*^+/rtTA^ from P14 to P35. (**A**-**D**) AAV-*TRE*-YFP was stereotactically injected into the corpus callosum of *Sox10*^+/rtTA^ or *Plp*-tTA mice at P14 or P35. Brain sections obtained 1 (AAV-1d) or 2 (AAV-2d) days after virus injection were co-immunostained by antibodies to YFP and Olig2, or by CC1 antibody and antibody to YFP, or by antibodies to YFP and NG2 (or PDGFRα). Shown are representative images for indicated mice at P14 or P35. *Sox10*^+/rtTA^ mice were fed with Dox for 3 days before stereotactic injection of the virus, while *Plp*-tTA mice had no Dox treatment. (**E** and **F**) Distributions of infected cell, as shown by co-immunostaining of YFP and TCF4, in the corpus callosum of *Sox10*^+/rtTA^ (E) or *Plp*-tTA (F) mice at P35. Note that infected cells in *Sox10*^+/rtTA^ mice stringently distributed within TCF4^+^ cell clustered region, whereas those in *Plp*-tTA mice distributed broadly in the corpus callosum. Solid arrows, regions with clustered TCF4^+^ cells; Open arrows, regions with fewer TCF4^+^ cells.

**Figure 3-figure supplement 1.**
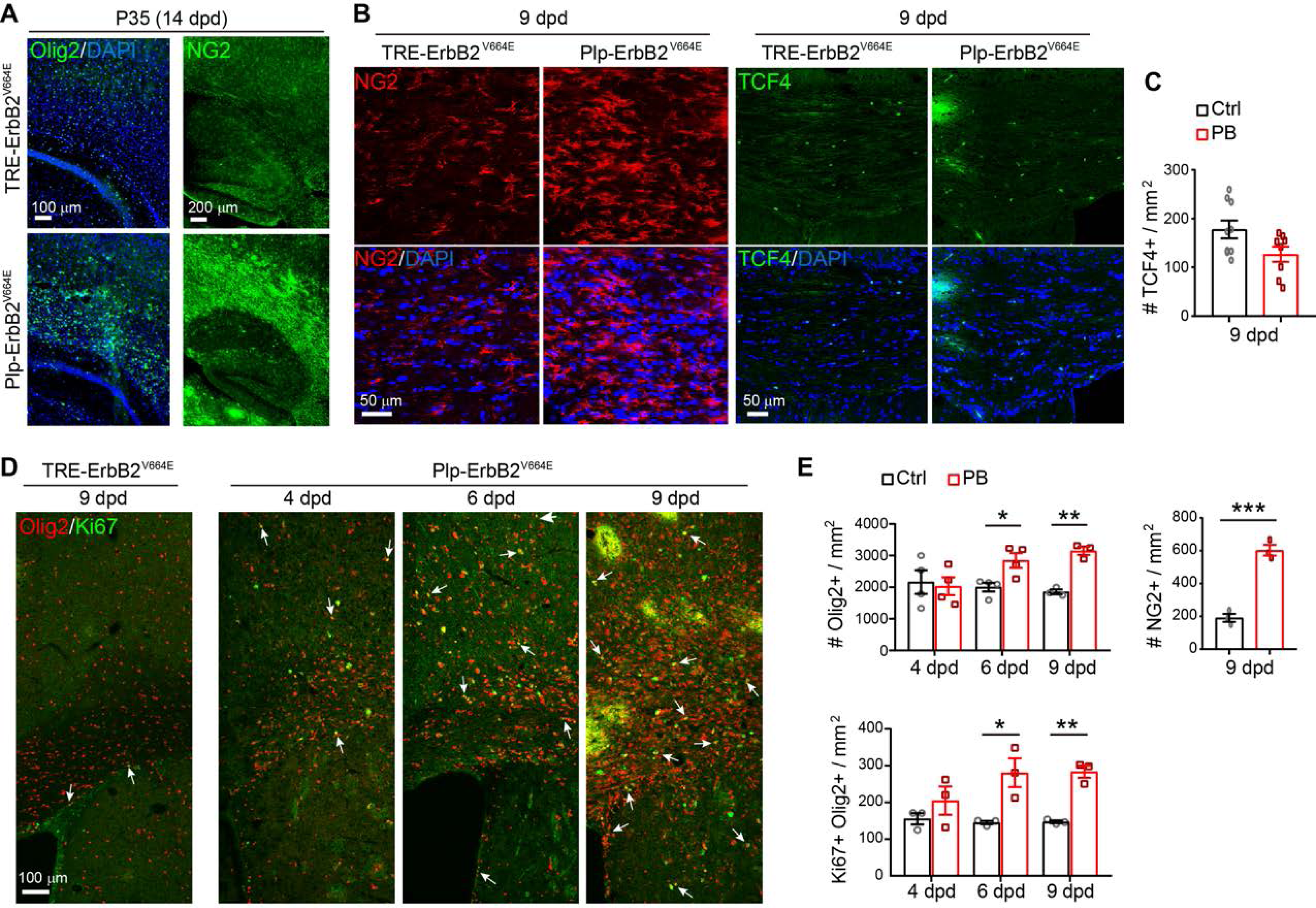
Pathological OL proliferation in subcortical white matter of *Plp*-ErbB2^V664E^ mice. (**A**) Dramatically increased Olig2^+^ and NG2^+^ cells in the subcortical white matter of *Plp*-ErbB2^V664E^ mice at 14 dpd. Sagittal sections of *Plp*-ErbB2^V664E^ and littermate control mice were immunostained by antibodies for Olig2 or NG2. (**B** and **D**) Immunostaining results of NG2 and TCF4 (B), Olig2 with Ki67 (D), in the corpus callosum of indicated mice. (**C** and **E**) Quantitative data of immunostaining results in *Plp*-ErbB2^V664E^ (PB) and control mice (Ctrl) with indicated Dox treatment were present as mean ± s.e.m., and analyzed by two-tailed unpaired *t* test. For TCF4^+^ density at 9 dpd, *t*_(15)_ = 2.1, *P* = 0.053. For Olig2^+^ density: at 4 dpd, *t*_(6)_ = 0.2923, *P* = 0.780; at 6 dpd, *t*_(6)_ = 3.16, *P* = 0.0196; at 9 dpd, *t*_(4)_ = 8.563, *P* = 0.001. For NG2^+^ density at 9 dpd, *t*_(4)_ = 9.912, *P* = 0.0006. For Olig2^+^Ki67^+^ density: at 4 dpd, *t*_(4)_ = 1.187, *P* = 0.301; at 6 dpd, *t*_(4)_ = 3.428, *P* = 0.027; at 9 dpd, *t*_(4)_ = 8, *P* = 0.0013.

**Figure 4-figure supplement 1.**
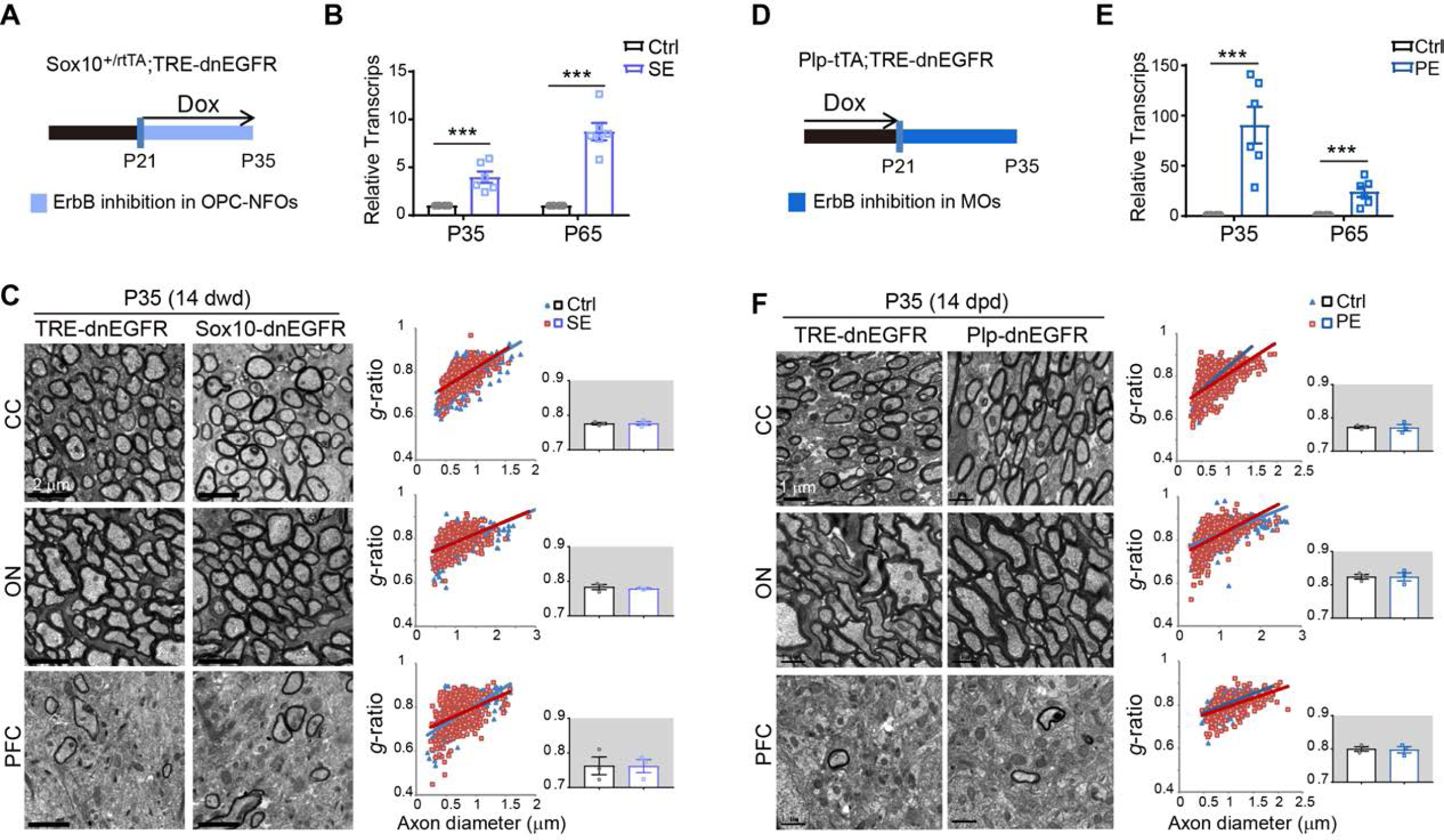
No myelin alteration in *Sox10*-dnEGFR (SE) or *Plp*-dnEGFR (PE) mice at P35 with 14 days of Dox treatment. (**A** and **D**) Dox treatment setting for indicated mice and littermate controls. (**B** and **E**) Without an antibody recognizing dnEGFR specifically, we examined the expression of dnEGFR/EGFR by real-time RT-PCR. As *Sox10*^+/rtTA^ targets a transient cellular stage, transcripts of dnEGFR/EGFR in the subcortical white matter of *Sox10*-dnEGFR mice were only 4- and 9-fold more than that of the endogenous EGFR in littermate controls (Ctrl) at P35 and P65, respectively, while that of *Plp*-dnEGFR mice were 90- and 24-fold more than the endogenous EGFR transcripts in littermate controls at P35 and P65, respectively. Data were presented as mean ± s.e.m., and analyzed by two-tailed unpaired *t* test. For *Sox10*-dnEGFR at P35, *t*_(10)_ = 5.044, *P* = 0.0005; at P65, *t*_(10)_ = 8.531, *P* < 0.0001. For *Plp*-dnEGFR at P35, *t*_(10)_ = 4.908, *P* = 0.0006; at P65, *t*_(10)_ = 4.515, *P* = 0.0011. (**C** and **F**) EM images of the corpus callosum (CC), optic nerve (ON), and prefrontal cortex (PFC) from *Sox10*-dnEGFR and littermate controls at P35 with 14 dwd, or from *Plp*-dnEGFR and littermate controls at P35 with 14 dpd. *g*-ratio was calculated for myelinated axons. Averaged *g*-ratio for each mouse (inset) were presented as mean ± s.e.m., and analyzed by two-tailed unpaired *t* test. In C, for CC, *t*_(4)_ = 0.1013, *P* = 0.924; for ON, *t*_(4)_ = 0.6191, *P* = 0.569; for PFC, *t*_(4)_ = 0.02485, *P* = 0.981. In F, for CC, *t*_(4)_ = 0.1443, *P* = 0.892; for ON, *t*_(4)_ = 0.01551, *P* = 0.988; for PFC, *t*_(4)_ = 0.1573, *P* = 0.883.

**Figure 5-figure supplement 1.**
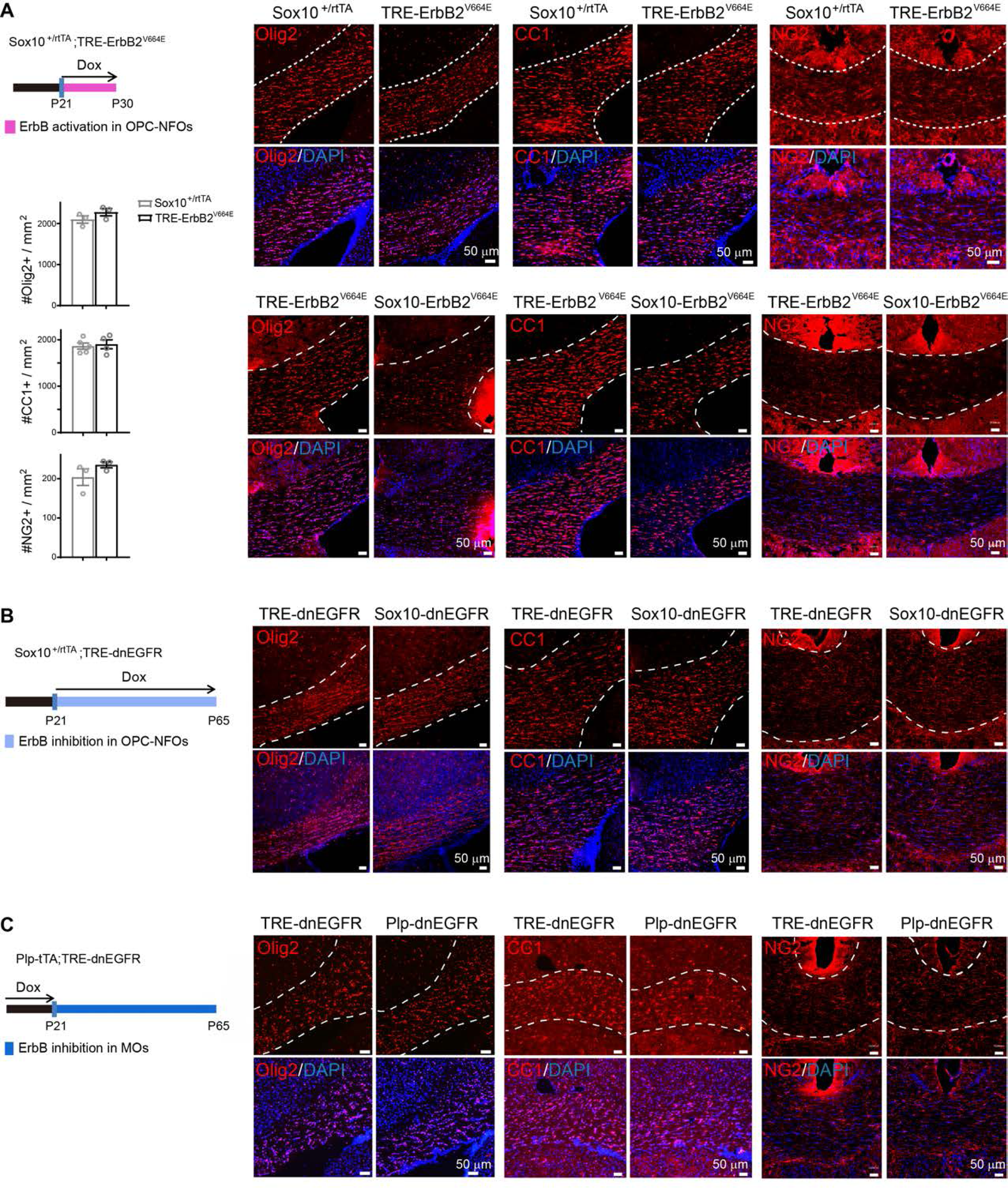
ErbB inhibition in OPC-NFOs increases OL numbers. (**A**-**C**) Olig2^+^, CC1^+^, and NG2^+^ cells in the corpus callosum of indicated mice at indicated ages were examined by immunostaining. Statistic results in (A) showed Olig2^+^, CC1^+^, and NG2^+^ cell densities were similar in the corpus callosum of *Sox10*^+/rtTA^ and littermate *TRE*-ErbB2^V664E^ mice at P30 (9 dwd). Data were from immunostaining of 3 mice for each group, presented as mean ± s.e.m., and analyzed by two-tailed unpaired *t* test. For Olig2^+^, *t*_(4)_ = 1.418, *P* = 0.229; for CC1^+^, *t*_(7)_ = 0.3431, *P* = 0.742; for NG2^+^, *t*_(4)_ = 1.394, *P* = 0.236.

**Figure 5-figure supplement 2.**
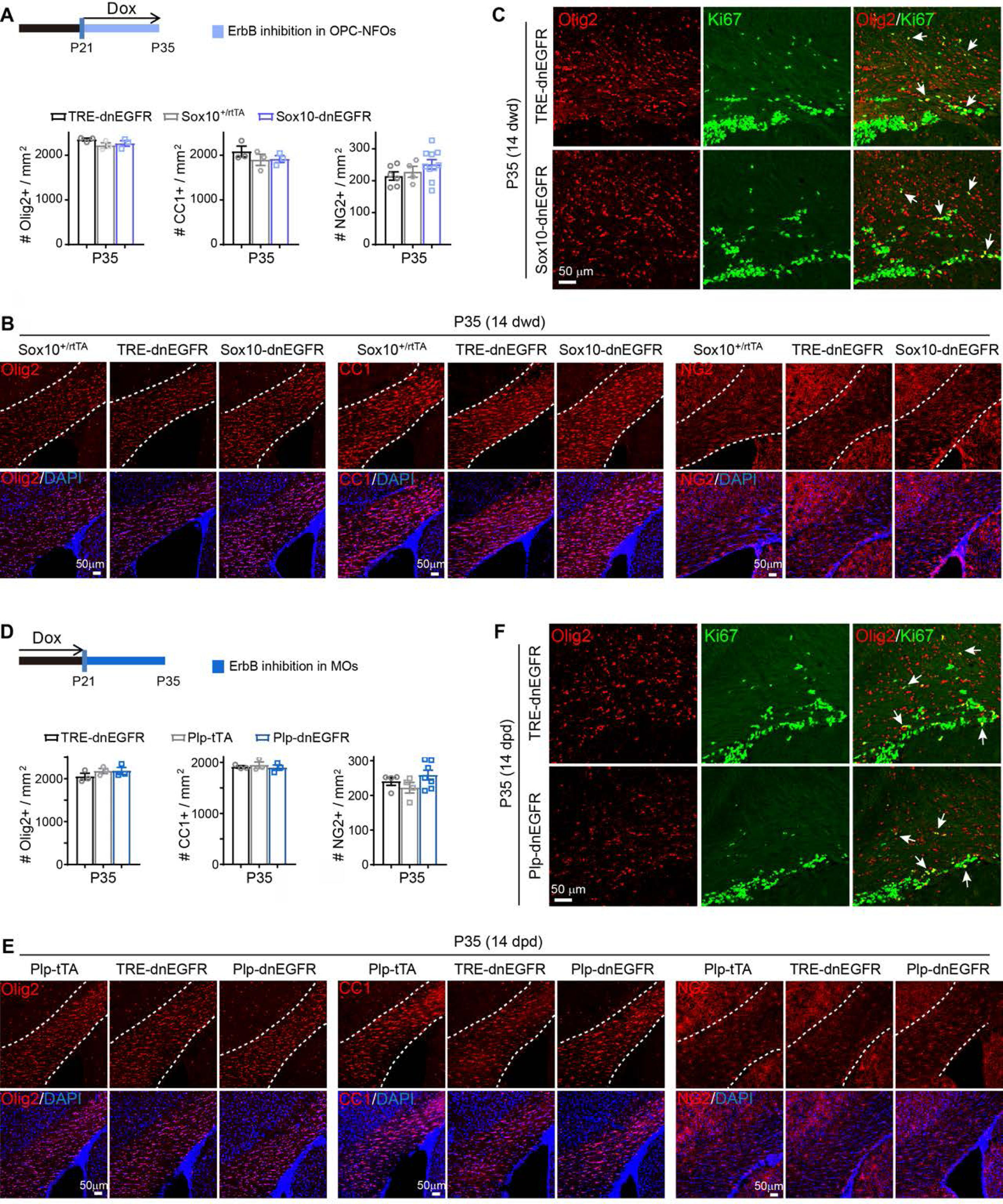
No changes in OL numbers or proliferation in *Sox10*-dnEGFR or *Plp*-dnEGFR mice at P35 with 14 days of Dox treatment. (**A** and **D**) Statistic results of Olig2^+^, CC1^+^, and NG2^+^ cell densities in the corpus callosum of indicated mice at P35. Data were from repeated immunostaining of 3 mice for each group, presented as mean ± s.e.m., and analyzed by one-way ANOVA. In A, for Olig2^+^, *F*_(2,6)_ = 1.651, *P* = 0.268; for CC1^+^, *F*_(2,6)_ = 0.8605, *P* = 0.469; for NG2^+^, *F*_(2,17)_ = 1.624, *P* = 0.226. In D, for Olig2^+^, *F*_(2,6)_ = 1.054, *P* = 0.405; for CC1^+^, *F*_(2,6)_ = 0.2694, *P* = 0.773; for NG2^+^, *F*_(2,12)_ = 1.633, *P* = 0.236. (**B** and **E**) Immunostaining results of Olig2^+^, CC1^+^, and NG2^+^ cells in the corpus callosum of indicated mice at P35. (**C** and **F**) Double immunostaining results of Olig2 and Ki67 in the corpus callosum of indicated mice at P35. Arrows, representative double positive nuclei.

**Figure 7-figure supplement 1.**
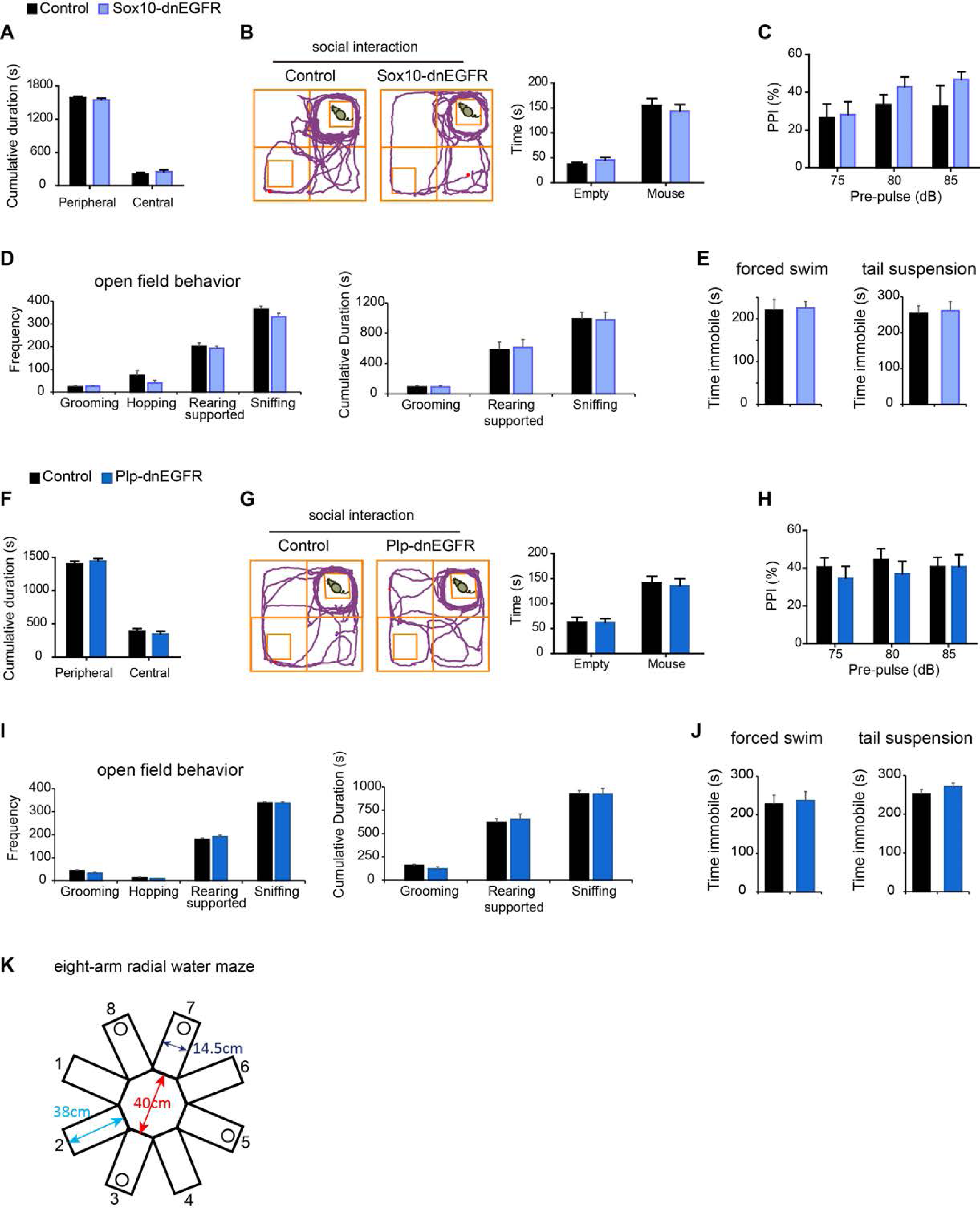
No behavioral abnormalities in sensory gating, social interaction, or mood behaviors revealed for *Sox10*-dnEGFR or *Plp*-dnEGFR mice. Behavioral performance of adult *Sox10*-dnEGFR mice with littermate controls (A-E), or *Plp*-dnEGFR mice with littermate controls (F-J), were tested in the open field test (**A** and **F**), social interaction test (**B** and **G**), prepulse inhibition (PPI) test (**C** and **H**), stereotyped behaviors in the open field (**D** and **I**), and the forced swim test and the tail suspension test (**E** and **J**). For zone analysis of open field tests, n = 13 mice for *Sox10*-dnEGFR and n = 11 mice for controls (two-way ANOVA test, *F*_(1, 44)_ = 0, *P* > 0.9999), while n = 14 mice for *Plp*-dnEGFR and n = 19 mice for controls (two-way ANOVA test, *F*_(1, 62)_ = 0.00017, *P* = 0.989). For PPI tests, n = 12 mice for *Sox10*-dnEGFR and n = 10 mice for controls (two-way ANOVA test, *F*_(1, 60)_ = 2.36, *P* = 0.13), while n = 12 mice for *Plp*-dnEGFR and n = 14 mice for controls (two-way ANOVA test, *F*_(1, 72)_ = 0.9134, *P* = 0.342). For social interaction tests, n = 13 mice for *Sox10*-dnEGFR and n = 12 mice for controls (two-way ANOVA test, *F*_(1, 46)_ = 0.027, *P* = 0.87), while n = 13 mice for *Plp*-dnEGFR and n = 14 mice for controls (two-way ANOVA test, *F*_(1, 50)_ = 0.1023, *P* = 0.75). For forced swim and tail suspension tests, n = 13 mice for *Sox10*-dnEGFR and n = 10 mice for controls (two-tailed unpaired *t* test, for forced swim, *t_(_*_21)_ = 0.1799, *P* = 0.859; for tail suspension, *t_(_*_21)_ = 0.2576, *P* = 0.799), while n = 13 mice for *Plp*-dnEGFR and n = 13 mice for controls (two-tailed unpaired *t* test, for forced swim, *t_(_*_24)_ = 0.2676, *P* = 0.791; for tail suspension, *t_(_*_24)_ = 1.189, *P* = 0.246). Data were presented as mean ± s.e.m.. Note that only male mice were used for PPI, social interaction, forced swim and tail suspension tests. (**K**) Illustration showing the setting for eight-arm radial water maze. Four hidden platforms were placed at the end of a same set of arms with 38-cm distance to the central zone at the training and test days. Mice started swimming with face to the arm end from No.1 arm in each trial, and the visited platform was removed before the next trial after 30-sec gap.

## Other Supplemental Files

**Figure 7-Video 1.** The performances recorded for *Sox10*-dnEGFR mice and *Plp*-dnEGFR mice, as well as their controls, in the 4th trial of eight arm radial water maze at the test day.

**Figure 6-Source data 1.** The Excel file contains the processed RNA-seq results of genes with differential expression in white matter tissues between *Sox10*-ErbB2^V664E^ mice and littermate *TRE*-ErbB2^V664E^ mice at P30 with 9 dwd.

**Figure 6-Source data 2.** The Excel file contains the processed RNA-seq results of genes with differential expression in white matter tissues between *Sox10*-dnEGFR mice and littermate *TRE*-dnEGFR mice at P35 with 14 dwd.

**Source data for graphs.** The zip file includes all raw numerical data in Prism files for graphs in each figure.

